# bHLH35 mediates specificity in plant responses to multiple stress conditions

**DOI:** 10.1101/2025.05.07.652729

**Authors:** Ranjita Sinha, Sara I. Zandalinas, María Ángeles Peláez-Vico, Abdul Ghani, Mather A. Khan, Sai Preethi Induri, Ahmad Bereimipour, Tara Ghandour, Andrew Ogden, Shao-Shan Carol Huang, Rajeev K Azad, David Mendoza-Cózatl, Trupti Joshi, Felix B. Fritschi, Ron Mittler

## Abstract

How biological systems respond to stress is a fundamental question in biology, primarily addressed using the reductionist approach of applying one stress condition at a time. In nature, however, organisms experience a multitude of stresses, simultaneously or sequentially, questioning the validity of the reductionist approach for predicting plant responses to stress under natural conditions. Here, we reveal that in the flowering plant *Arabidopsis thaliana,* the transcriptional regulator bHLH35 is required for plant survival under a specific set of stress conditions that includes a combination of salinity, excess light, and heat, occurring simultaneously (but not for each of these stresses applied individually or in any other combination). Under these conditions, bHLH35 interacts with NAC069 and binds the promoter of *LBD31*, also specifically required for survival under the 3-stress combination. Our findings uncover a high degree of specificity in the response of organisms to stress, a specificity that would not have been revealed using the reductionist approach, and one that should be taken into consideration when developing agronomically important crops with heightened resilience to climate change.

## Main Text

Global warming and climate change are altering our environment, negatively impacting the growth and yield of crops, and threatening food security^1–13^. Especially notable are the elevated day and night temperatures, and the increase in the frequency, duration and intensity of weather events, such as heat waves, floods, and droughts. In many instances, some of these weather events occur simultaneously or sequentially, subjecting plants to conditions of ‘stress combination’^1–4,14^. For example, when heat waves occur during periods of drought, when drought, salinity, and heat stresses combine, or during cycles of drought and waterlogging stresses^1–3,5,14,15^. While previous studies determined the responses of plants to different abiotic stresses, such as heat, salinity, or drought, applied individually (the ‘reductionist’ approach), much less is known about plant responses to different combinations of these conditions (a ‘holistic’ approach that takes into consideration the high complexity of environmental stress conditions occurring in nature). This lack of knowledge is likely to severely hinder attempts to develop agronomically important crops with heightened resilience to climate change and global warming, essential for our survival in the coming years^2,5,9,12^.

To determine the role of different plant pathways regulating responses to compound stress conditions, we mined the large dataset of *Arabidopsis thaliana* (*A. thaliana*) transcriptomic responses to stress (https://www.ncbi.nlm.nih.gov/geo/), with a special focus on conditions of multifactorial stress combination^4,16^. We identified over 130 transcripts, including 12 that encode transcription factors (TFs), significantly altered in their expression in response to combinations of 2-, 3-, 4-, 5-, and 6-different abiotic stress conditions, applied simultaneously (Figs. 1a, S1). Of these, we focused on bHLH35^17^ (At5g57150), as the function of this TF is largely unknown in *A. thaliana*, suggesting that it has no identified role in responses to different environmental stress conditions applied separately. To functionally test the role of bHLH35 in different stress combinations, we subjected wild type (WT) and two independent knockout alleles of *bHLH35* (*bhlh35‗1* and *bhlh35‗2*; Fig. 1b; Table S1) to five different abiotic stresses in all possible combinations (Fig. 1b; Table S2). Surprisingly, *bhlh35* mutants were exclusively impaired in their survival under a combination of salt (S), excess light (EL), and heat stress (HS), or any other combination of stresses that included these 3 stresses (S+EL+HS; Fig. 1b). The unique susceptibility of *bhlh35* mutants to S+EL+HS (Fig. 1b) occurred under a range of S, EL, and HS conditions (Fig. 1c, Table S2), underscoring the specificity of bHLH35 to this stress combination. Using the highest settings of S+EL+HS, we also demonstrate that *bhlh35‗2* could be genetically complemented with the *bHLH35* cDNA expressed under the control of its native promoter (Fig. 1d). In addition, we show that overexpressing the *bHLH35* cDNA using the *CaMV35S* promoter specifically enhances tolerance of *A. thaliana* to the S+EL+HS combination (Fig. 1e). The susceptibility of the *bhlh35* mutants to S+EL+HS was also observed in a simulated field environment (Figs. 1f, Table S2), highlighting the importance of bHLH35 for plant survival in nature. In support of the simulated field experiments, *A. thaliana* ecotypes with high expression level of *bHLH35* were more resilient to conditions of S+EL+HS, compared to ecotypes with low expression level^18^ (Fig. 1g), and single nucleotide polymorphism (SNPs) in the *bHLH35* gene correlated with high irradiance, heat stress, and ozone levels in a geoclimatic variable associations of 879 *A. thaliana* accessions^19^ (Fig. 1h). These results support a role for bHLH35 under natural conditions.

**Fig. 1.**
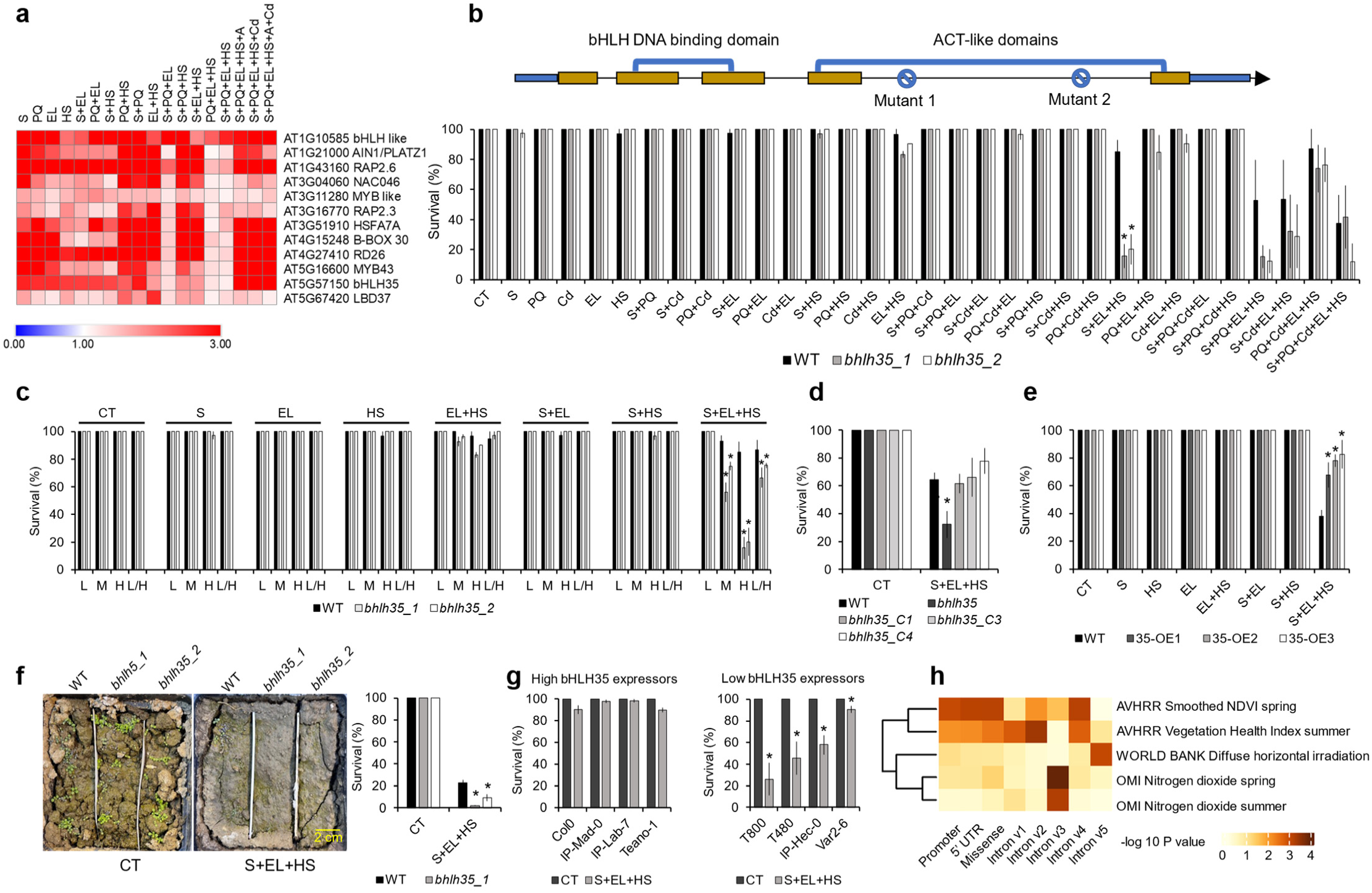
The transcription factor bHLH35 is required for plant survival under conditions of stress combination. **a,** *Arabidopsis* transcription factors (TFs) transcripts expressed in response to different stress combinations. **b,** Map of the *bHLH35* gene with the position of T-DNA insertions in the two independent *bhlh35* mutants (Top), and survival of wild type (WT) and the two independent *bhlh35* mutants (Table S1) under control (CT) and stress combination conditions (salt, S; the herbicide paraquat, PQ; cadmium, Cd; excess light, EL; and heat stress, HS; in all possible combinations; Middle). **c**, Survival of WT and the two *bhlh35* mutants subjected to CT, S, HS, and/or EL at different stress levels (L, low; M, medium; H, high; L/H, low S high HS and EL). **d**, Complementation of *bhlh35_2* with the *bHLH35* cDNA expressed under its native promoter (C, complementation). **e**, Survival of WT and three *bHLH35* gain-of-function lines (cDNA expressed under the control of the *CaMV35S* promoter) mutants subjected to CT, S, HS, and/or EL conditions. **f**, Survival of WT and the two *bhlh35* mutants subjected to CT or S+EL+HS under simulated field conditions. **g**, Survival of different *A. thaliana* ecotypes with high (Left) or low (Right) expression level of *bHLH35* under CT or S+EL+HS conditions. **h**, Correlation between single nucleotide polymorphism (SNPs) in the *bHLH35* gene and environmental conditions obtained from the GenoCLIM v2.0 database of geoclimatic variable associations for 879 *A. thaliana* accessions. Significant associations were found with high irradiance, heat stress, and ozone. *Two-tailed Student’s t-test (p ≤ 0.05).

Focusing on the combination of S+EL+HS, we conducted a transcriptomics analysis of WT and one of these mutants (*bhlh35‗2*; referred herein as *bhlh35*) subjected to S, EL, and HS, in all possible combinations (Fig. S2a; Tables S1, S2). This analysis revealed marked differences between the transcriptional responses of WT and *bhlh35* to the different stress conditions (Figs. 2a, S2b, S2c; Tables S3-S5). Comparing the transcriptomics responses of WT to that of *bhlh35* under each of the different stress conditions applied revealed that only 10 different transcripts were common to all *bHLH35*-dependent responses of WT (indicated in bold in Fig. 2a) to each of these stress conditions (Arrow in Fig. 2b; Table S6). This finding suggested that bHLH35 could play different roles under different stress conditions. Considering bHLH35 specificity (Figs. 1, 2, S2; Tables S3-S6), we focused on the combination of S+EL+HS and defined two classes of transcripts: Transcripts that were altered in their expression in WT specifically under the S+EL+HS combination but did not respond at all in *bhlh35* (Class 1; Fig. 2c; Table S7), and transcripts that were significantly altered in their expression in *bhlh35* in response to the 3-stress combination but did not respond at all in WT (Class 2; Fig. 2c; Table S8). While Class 1 transcripts were enriched in ethylene- and biotic stress-response transcripts, Class 2 transcripts were enriched in development- and general stress-response transcripts (Figs. 2d, S3). Analysis of mutants deficient in ethylene signaling (Table S1) revealed however that these mutants had altered survival under conditions of EL+HS as well as S+EL+HS (Fig. 2e). We next identified transcripts that had a bHLH binding domain (G- and E-boxes) in their promoters from each of these two groups (Fig. 2c) and conducted a cluster analysis to identify transcripts unique to the combination of S+EL+HS in each group (Figs. 2f, S4).

**Fig. 2.**
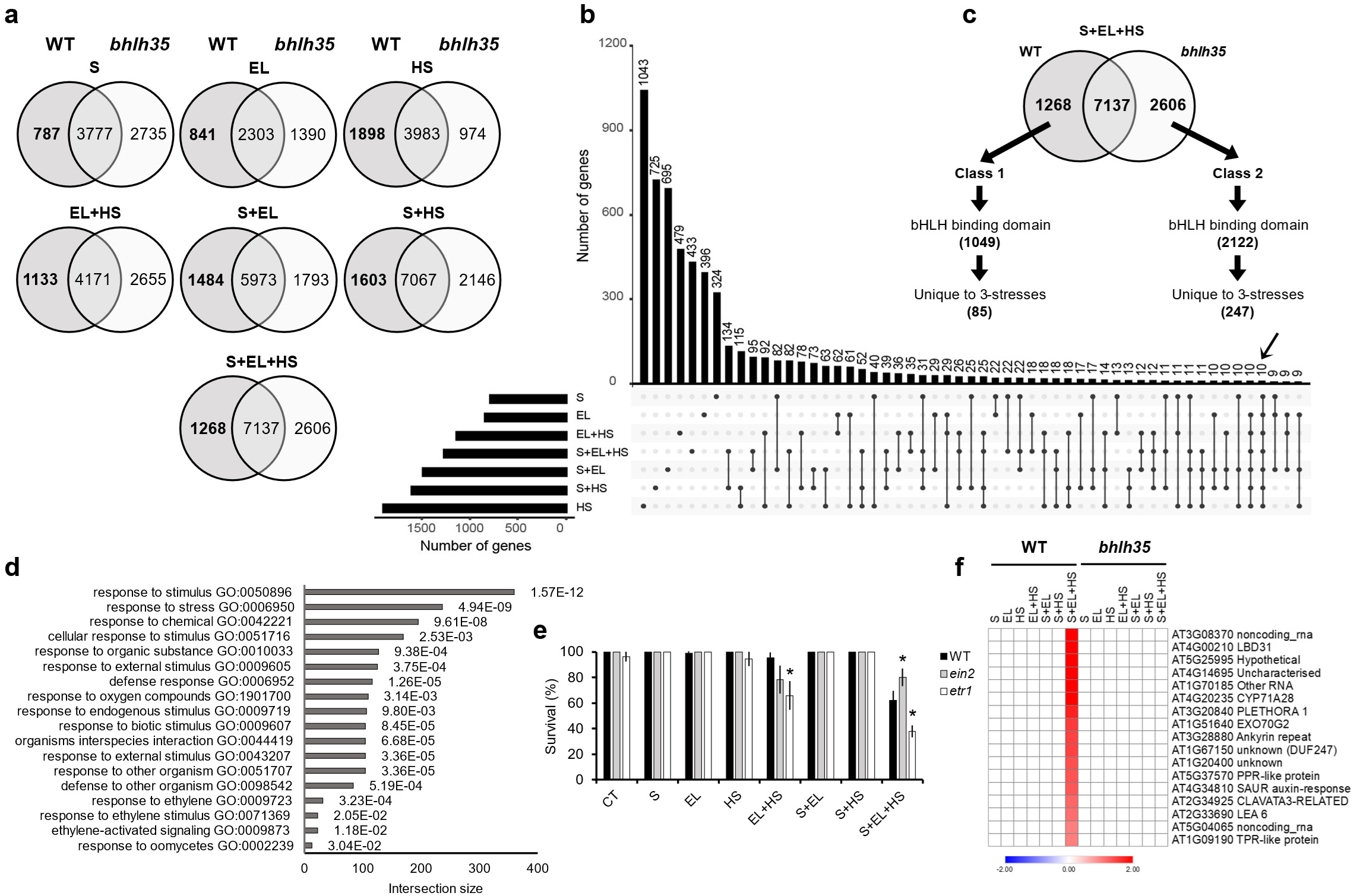
bHLH35 is required for the expression of stress- and ethylene-response transcripts under a combination of salt, excess light, and heat stress. **a**, Venn diagrams showing the overlap between transcripts significantly altered in their expression in WT and *bhlh35_2* in response to salt (S), excess light (EL), and heat stress (HS) in all possible combinations. **b**, Upset plot showing the overlap between all *bHLH35*-dependent transcripts expressed in WT under the different stress conditions (From a, bold). Arrow indicates transcripts common to all *bHLH35*-dependent transcripts expressed in WT under the different stress conditions. **c**, The proportion of transcripts encoded by genes with a bHLH DNA-binding domain in their promoter, and the proportion of the latter with a unique expression pattern to S+EL+HS (Tables S7, S8) among the *bHLH35*-dependent transcripts expressed in WT or *bhlh35_2* under a combination of S+EL+HS. **d**, Gene ontology (GO) analysis of *bHLH35*-dependent transcripts (1,268 from Class 1, c) identifying stress- and ethylene-response transcripts (GO analysis of the 2,606 transcripts from Class 2 is shown in Fig. S3). **e**, Altered survival of ethylene biosynthesis and sensing mutants (Table S1) under conditions of EL+HS and S+EL+HS. **f**, Heatmap showing a partial list of a cluster of *bHLH35*-dependent transcripts that are specifically expressed in wild type (WT) under conditions of S+EL+HS (complete list is shown in Fig. S4a). *Two-tailed Student’s t-test (p ≤ 0.05), comparing WT to ethylene mutants within each treatment.

Analysis of mutants (two independent alleles) of randomly selected genes encoding some of the transcripts included in Class 1 and Class 2 (Figs. 3a, S5; total of 16 genes were tested with two independent mutants each; Table S1), identified *ERF111*^20^ and *RAP2.3*^21^, involved in ethylene signaling, *bHLH42*^22^ (*TT8*, involved in flavonoids biosynthesis), *ACHT4*^23^ a redox-response His-rich thioredoxin, and *LBD31*^24^ a TF involved in development and secondary metabolism, as essential for plant survival specifically under the combination of S+EL+HS (similar to *bhlh35*; Figs. 1b, 3b, S5). Of these, only *LBD31* was specifically expressed under the 3-stress combination conditions in WT but not in *bhlh35* (Fig. 3a). We next conducted a yeast 2-hybrid analysis of bHLH35 against an *A. thaliana* whole-genome TF library^25^ (Fig. 3c). This screen revealed that bHLH35 interacted with several other TFs (Figs. 3c, S6a). Functional analysis of some of these interactors showed that, although none of them had a unique expression pattern like *bHLH35* (Figs. 3a, S6b), mutants of *MYB12*^26^ and *NAC069*^27^, displayed a similar survival phenotype to that of *bhlh35*, whereas mutants of *MYC2*^28^ were specifically required for survival under the combinations of EL+HS as well as S+EL+HS (Figs. 1, 3d; Table S1). As LBD31 was the only TF with an expression pattern (Figs. 2f, 3a), as well as a survival phenotype (Fig. 3b), unique to the 3-stress combination, we focused on this TF and conducted a targeted yeast 1-hybrid (Y1H) and electrophoretic mobility shift assay (EMSA) analyses to determine if bHLH35 binds to its promoter. While in the Y1H assay bHLH35 showed binding to the *LBD31* promoter (Fig. 4a), in the EMSA study it needed to interact with NAC069 before it could bind the *LBD31* promoter (Fig. 4b). Gene regulatory network (GRN) analyses^29,30^ of *bHLH35* function further identified flavonoids, abscisic acid (ABA), as well as heat shock transcription factors (HSFs) and other stress response transcripts, associated with bHLH35 function (Figs. 4c, 4d, S7, S8; Table S9). As transcripts involved in flavonoid metabolism were identified by our GRN analyses (Figs. 4c, S8), *bHLH42*^22^ (*TT8*, a known regulator of flavonoid biosynthesis) was essential for plant survival under the combination of S+EL+HS (Fig. 3b), and homologs of bHLH35 and LBD proteins were previously associated with flavonoid metabolism in different plants^31–34^, we tested whether augmenting flavonoid metabolism by treatment of plants with naringenin^35,36^ could rescue the *lbd31* mutants. Indeed, naringenin application could rescue the *lbd31* phenotype specifically under the 3-stress combination (Fig. 4e, top). In contrast, naringenin could not rescue the *rap2.3* mutants involved in ethylene signaling under the 3-stress combination (Fig. 4e, bottom). Under conditions of S+EL+HS, bHLH35 therefore regulated a unique set of transcripts that was specific to the 3-stress combination condition and associated with flavonoid biosynthesis and ethylene signaling pathways, required for plant survival (Fig. 4f).

**Fig. 3.**
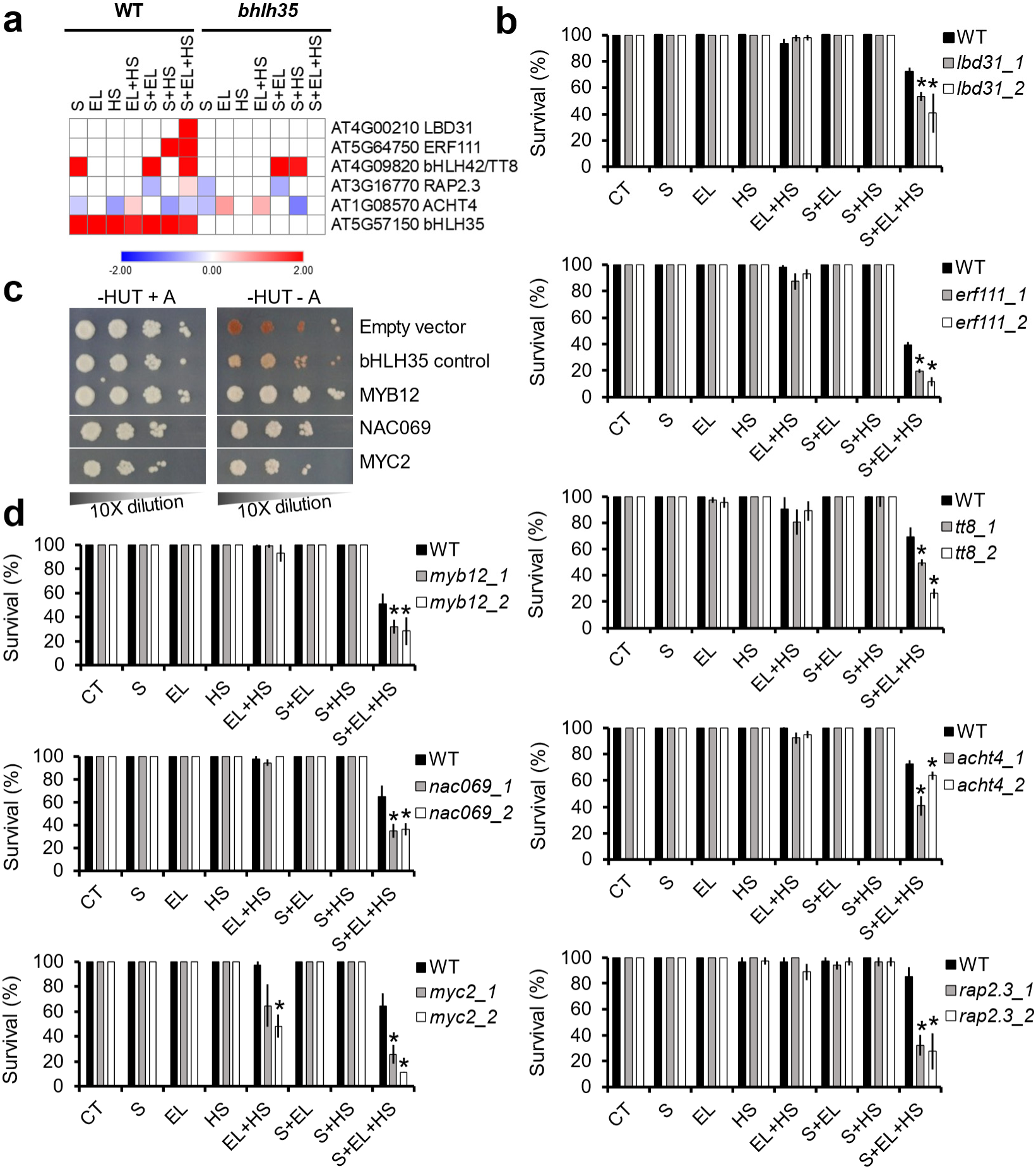
The expression of different *bHLH35*-dependent transcripts and bHLH35 interactors is specifically required for survival under a combination of salt, excess light, and heat stress. **a,** Expression of *bHLH35* and transcripts that depend on *bHLH35* for expression under a combination of salt (S), excess light (EL), and heat stress (HS). **b**, Two independent mutants for each of the genes encoding transcripts that require *bHLH35* for expression under conditions of S+EL+HS (From a; Table S1) are deficient in survival specifically under this set of stress combination conditions (Specific survival of *bhlh35* under conditions of S+EL+HS is shown in Fig. 1b; Survival of other *bHLH35*-dependent transcripts is shown in Fig. S5b). **c**, Selected yeast 2-hybrid results for transcription factors (TFs) that interact with bHLH35 (Full results are shown in Fig. S6a). **d**, Survival of two independent mutants for each of the TFs shown in c to interact with bHLH35 under conditions of S, EL, and HS in all possible combinations (Table S1). *Two-tailed Student’s t-test (p ≤ 0.05), comparing WT to the different mutants in genes with *bHLH35*-dependent expression within each treatment.

**Fig. 4.**
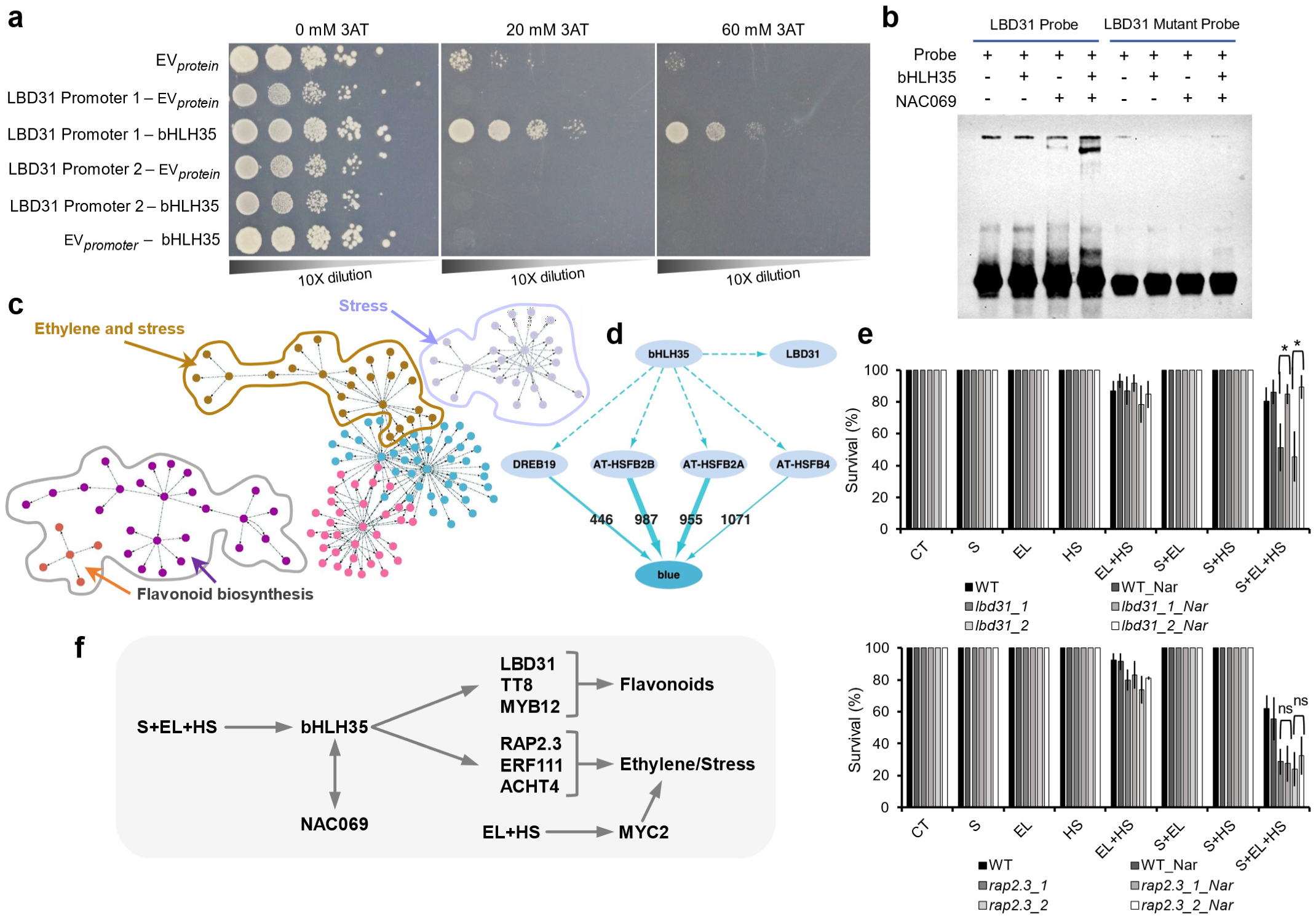
bHLH35 interacts with NAC069 and binds the promoter of *LBD31*, driving both ethylene- and flavonoid-associated pathways during a combination of salt, excess light, and heat stress. **a,** Yeast 1-hybrid analysis showing that bHLH35 binds the promoter (LBD31 promoter 1 fragment) of *LBD31* (EV, empty vector; Table S10). **b**, Gel mobility shift assay showing that the *in vitro* binding of bHLH35 to the promoter 1 fragment of *LBD31* requires NAC069. **c**, Gene regulatory network (GRN) analysis of *bHLH35* unravels links to stress, flavonoid-, and ethylene-associated transcripts (Table S9). **d**, A sub module in a weighted gene co-expression network analysis (WGCNA) of bHLH35 that regulates responses to heat stress (the blue module; Fig. S8, Table S11). Dashed arrows: Co-expression and promoter binding; Solid arrows: Co-expression and significant motif enrichment. **e**, The flavonoid naringenin (Nar) rescues the survival phenotype of *lbd31*, but not *rap2.3* mutants, under a combination of S+EL+HS. **f**, A model for the function of bHLH35. bHLH35 is shown to be required for the expression of *LBD31*, *bHLH42* (*TT8*), and *MYB12*, all associated with flavonoids metabolism and specifically required for the survival of *Arabidopsis thaliana* seedlings under conditions of S+EL+HS combination. *bHLH35* is also shown to be required for the expression of *RAP2.3*, *ERF111*, and *ACHT4*, all associated with ethylene and stress metabolism and required for survival under conditions of S+EL+HS combinations. *Two-tailed Student’s t-test (p ≤ 0.05).

Taken together, our findings demonstrate an important principle in biology, *i.e.,* that different combinations of different environmental stresses require different genetic programs for survival. The survival of *A. thaliana* seedlings under conditions of S+EL+HS is shown here to require a specific set of TFs that controls a specific set of transcripts that are not required for survival under S, EL, or HS in all possible combinations, other than S+EL+HS. We further show that a distinct combination of different pathways is required for survival under each condition of stress combination. Thus, *bHLH35* regulated ethylene signaling pathways required for EL+HS acclimation (together with MYC2), as well as flavonoid metabolism and ethylene signaling pathways required for S+EL+HS (together with NAC069; Fig. 4f). Such specificity suggests that different organisms utilize a genetically encoded ‘stress combination-specific response’, or responses, which could only be revealed using a ‘holistic’ approach of testing all possible stress conditions in different combinations. In the context of global warming, climate change, and increased pollution levels, this principle is also important, as humans, microbiomes, animals, and plants are continuously subjected to a multitude of different stress conditions in an environment that continues to increase in its complexity with each year^4,11,37,38^. Studies of multifactorial stress combination, or global change factor combinations, have recently revealed that with the increasing complexity of stress conditions impacting a plant, microbiome, or ecosystem, their functions and services dramatically decline^4,16,39–44^. A significant shift in the way we conduct experiments (towards a more ‘holistic’ approach) is therefore needed to address our future challenges as a society on this planet.

## Acknowledgments

We thank the Arabidopsis Biological Resource Center (https://abrc.osu.edu/) for seeds of Arabidopsis mutants used in this study. This work was supported in part through the NYU IT High Performance Computing resources, services, and staff expertise.

## Funding

National Science Foundation IOS-2414183 (RM, FBF)

National Science Foundation IOS-2110017 (RM, FBF, TJ)

National Science Foundation IOS-2343815 (RM, TJ)

National Science Foundation MCB-2224839 (DMC, RM)

National Institutes of Health R35GM138143 (SCH)

## Author contributions

Conceptualization: RM, SIZ, FBF

Methodology: RS, MAPV, AG, MAK, SPI, AB, AO, TG, SCH

Investigation: RM, SIZ, FBF, RS, MAPV, MAK, DMC, TJ, FBF, RKA, TG, SCH

Visualization: RS, MAPV, RM

Funding acquisition: RM, FBF, TJ, DMC

Project administration: RM

Supervision: RM, DMC, TJ, RKA, SCH

Writing – original draft: RM, RS, MAPV, SIZ

Writing – review & editing: RM, SIZ, RS, MAPV, DMC, SCH

## Data and materials availability

All materials used in the analysis are available upon request from the Corresponding Author. All data are available in the main text or the supplementary materials. RNA-Seq data is available at Gene Expression Omnibus (GEO) under accession GSE281968.

## Methods

### Plant growth, stress treatments, and mutant screening

*Arabidopsis thaliana* wild-type Col-0 (WT) and two independent loss-of-function mutants of *bHLH35* (AT5G57150; Table S1) were subjected to the following individual treatments and their different combinations on plates containing ½ Murashige and Skoog (½ MS; Caisson Labs, UT, USA; MSP09) medium, pH 5.8, with 1% phytagel (Table S2): CT (control; ½ MS, 21 °C, 50 µmol m^−2^ s^−1^, pH 5.8), Cd (cadmium; ½ MS, 21 °C, 50 µmol m^−2^ s^−1^, pH 5.8, 5 µM CdCl_2_), EL (excess light; ½ MS, 21 °C, pH 5.8, 700 µmol m^−2^ s^−1^), HS (heat stress; ½ MS, 50 µmol m^−2^ s^−1^, pH 5.8, 33 °C), S (salt stress; ½ MS, 21 °C, 50 µmol m^−2^ s^−1^, pH 5.8, 50 mM NaCl), and PQ (paraquat; ½ MS, 21 °C, 50 µmol m^−2^ s^−1^, pH 5.8, 0.05 µM paraquat)^16^ (Table S2). All other experiments comparing WT and *Arabidopsis* loss-of-function mutants (Table S1), or gene complementation lines for the *bhlh35* mutant were performed with S, EL and HS and all their combinations as described above and below (see also Table S2). Seeds of *A. thaliana* WT and mutants (Figs. 1-4, S5, Table S1) were sterilized with chlorine gas^16^ and placed on rectangular plates (12 cm width; Avantor, VWR Funding Inc, PA, USA; 75780-348) containing ½ MS supplemented with either salt, cadmium, or paraquat or combination of these based on the stress combination applied^16^ (Table S2). About 15–20 seeds of wild-type Col-0 and two independent homozygous genotypes of each mutant (Table S1) were placed side-by-side on the same plate and stratified at 4 °C overnight. Seedlings were grown vertically for 6 days (21 °C temperature, 50 µmol m^−2^ s^−1^) under continuous light conditions and subjected to heat (HS; 33 °C) and/or excess light (EL; 700 µmol m^−2^ s ^−1^) for 3 days before scoring for survival^16^. For abiotic stresses and their combinations not involving EL and/or HS, seeds were allowed to germinate and grow in the presence or absence of stress conditions for 9 days. Percent survival was measured for all plates at the same time (9 days)^16^. For experiments with different levels of S, EL, and/or HS*, Arabidopsis* WT and the two *bhlh35* mutants were screened for survival under low (L; 35mM NaCl, 31 °C HS, 500 µmol m^−2^ s^−1^), mild (M; 40 mM NaCl, 32 °C HS, 600 µmol m^−2^ s^− 1^), high (H; 50 mM NaCl, 33 °C HS, 700 µmol m^−2^ s^−1^), and low/high (L/H; 35mM NaCl, 34 °C HS, 800 µmol m^−2^ s^−1^) after 5, 7, and 10 days post stress application, as described in^16^. For Arabidopsis ecotype stress experiments, *A. thaliana* ecotypes (The 1001 Genomes Consortium; https://1001genomes.org^18,19^) with high or low steady-state *bHLH35* transcript expression levels^18^ (based on GSE80744) were screened for survival under CT and S+EL+HS (50 mM NaCl, 33 °C HS, 700 µmol m^−2^ s^−1^), as described above. For survival of Arabidopsis WT and the two *bhlh35* mutants under simulated field conditions (Fig. S9), seeds of the different lines were germinated side-by-side in plastic pots (10×10×10 cm) filled with field soil (Mexico silt loam; fine, smectitic, mesic Vertic Epiaqualf) obtained from the Bradford Research Center^45^ (Missouri Agricultural Experiment Station, Columbia, Missouri, USA, 38°53′N, 92°12′W). Soil for CT pots was watered with plain water, while soil for S+EL+HS was supplemented with 50 mM NaCl once at the beginning of the experiment. Seedlings were grown in a greenhouse (https://plantgrowthfacilities. missouri.edu/eastcampus.htm) under natural light shaded to 150 µmol m^−2^ s^−1^ at a temperature of ∼22 °C, day, ∼15 °C night. Five days post germination, CT non salt treated plants were kept under the conditions described above (CT conditions), while salt treated pots were subjected to EL and HS (for a combination of S+EL+HS), by removing them from shading, placing them under a heater (Kalglo®-Infrared Heater HS-2420 I 65” I 240V I 2000W, Fogelsville, PA, USA; Kimball, B. A. Theory and performance of an infrared heater for ecosystem warming. *Glob. Chang. Biol.* **11**, 2041-2056, 2005) and supplementing the natural daylight (500-600 µmol m^−2^ s^−1^) with LED lights (CRAFTSMAN 9000-Lumen LED twin light, model-CMXELAYMPL1029, CRAFTSMAN Mississauga, ON, CA; an additional 150-200 µmol m^−2^ s^−1^ above natural conditions), for a total of 650-800 µmol m^−2^ s^−1^ (Fig. S9). Four to five days post stress application, the survival of CT and S+EL+HS treated plants was determined as described by^16^. For survival experiment in the presence or absence of the flavonoid Naringenin^35^ (Indofine Chemical Company, NJ, USA; N-101), ½ MS plates with or without salt (50 mM) were supplemented with 100 µM Naringenin as described in^35^ and the experiment was conducted as described above (Table S1). Each treatment was repeated in at least three technical repeats and conducted in at least three biological replicates.

### RNA-Seq analysis

About 125-150 *A. thaliana* wild-type Col-0 and *bhlh35* (*bhlh35_2*) seedlings were grown horizontally for 6 days on separate sets of plates containing ½ MS with and without salt (50 mM NaCl) in three biological repeats. After 6 days, seedlings were subjected to the individual and combined stresses of S, EL and HS as described above except that EL and/or HS were applied for 1.5 hours before sampling. Whole seedlings for each stress treatments and control were sampled (flash freeze in liquid nitrogen) in three replicates after the 1.5 hours of EL and/or HS^16^. Total RNA was isolated using RNAeasy plant mini kit (Qiagen, MD, USA; 74904). RNA libraries for sequencing were prepared by Novogene Co. Ltd (https://en.novogene.com/website, Sacramento, CA) using standard Illumina protocols. RNA sequencing was performed using NovaSeq 6000 PE150 by Novogene Co. Ltd. Quality control was conducted using FastQC v0.11.9 (https://www.bioinformatics.babraham.ac.uk/projects/fastqc/), and summary reports were generated with MultiQC v1.12 (https://github.com/MultiQC/MultiQC/). Read trimming was performed to remove low-quality bases and adapter sequences using Trim Galore v0.6.4 (https://www.bioinformatics.babraham.ac.uk/projects/trim_galore/). Alignment of sequence reads to the Arabidopsis reference genome TAIR10, release 57 (https://ftp.ensemblgenomes.ebi.ac.uk/pub/plants/release-57/fasta/arabidopsis_thaliana/) was performed with Hisat2 v2.2.1 (Kim, D., Paggi, J. M., Park, C., Bennett, C. & Salzberg, S. L. Graph-based genome alignment and genotyping with HISAT2 and HISAT-genotype. *Nat. Biotechnol.* **37**, 907–915, 2019). Aligned sequences were sorted with Samtools v1.9 (Danecek, P. et al. Twelve years of SAMtools and BCFtools. Gigascience **10**, giab008, 2021) to facilitate downstream analyses. Transcript assembly and quantification were conducted using Cufflinks v2.2.1 (Trapnell, C. et al. Differential gene and transcript expression analysis of RNA-seq experiments with TopHat and Cufflinks. *Nat. Protoc.* **7**, 562–578, 2012) with the Ensembl annotation file from the same source as the reference genome. Principal Component Analysis (PCA) was performed to assess the clustering of biological replicates, confirming the reproducibility and consistency of the experimental conditions. Differential expression analysis was performed using Cuffdiff v2.2.1 (Trapnell, C. et al. Differential analysis of gene regulation at transcript resolution with RNA-seq. *Nat. Biotechnol.* **31**, 46–53, 2013) to identify differentially expressed genes (DEGs). Genes were considered differentially expressed if they had a q-value < 0.05. Functional annotation, quantification of overrepresented gene ontology (GO) terms (p < 0.05) and KEGG enrichment were conducted in g:Profiler (Raudvere, U. et al. g:Profiler: a web server for functional enrichment analysis and conversions of gene lists (2019 update) *Nucleic Acids Res.* **47**, W191–W198, 2019). The presence of bHLH G-box or E-box motifs in the promoter (500 bp upstream) of genes was performed in MEME Suit 5.5.4 (https://meme-suite.org/meme/index.html) using the FIMO motif scanning tool (Grant, C. E., Bailey, T. L., Noble, W. S. FIMO: Scanning for occurrences of a given motif. *Bioinformatics* **27**, 1017–1018, 2011). UpSet plots were generated using the UpSetR (Conway, J. R., Lex, A. & Gehlenborg, N. UpSetR: An R package for the visualization of intersecting sets and their properties. *Bioinformatics* **33**, 2938–2940, 2017) package. Heatmaps were generated using Morpheus (https://software.broadinstitute.org/morpheus/). Venn diagrams were created in VENNY 2.1 (BioinfoGP, CNB-CSIC; https://bioinfogp.cnb.csic.es/tools/venny/index.html). RNAseq data files are available at GEO (GSE281968).

### Gene regulatory network analysis

Gene Regulatory Network (GRN) analysis was performed using DIANE^29^ (Differential Inference Analysis of Expression). RNA-Seq data was normalized, and low-expression genes were filtered out, retaining genes with robust regulatory signals to infer gene-gene interactions based on differential expression profiles, leveraging both co-expression and causative regulatory evidence to capture direct and indirect regulatory relationships. DIANE parameters were optimized, with a significance threshold set to a p-value <0.05, and multiple testing was addressed through the Benjamini-Hochberg correction. The resultant output, an edge list of significant gene interactions, was prepared in Cytoscape 3.10 (Otasek, D., Morris, J. H., Bouças, J., Pico, A. R. & Demchak, B. Cytoscape Automation: Empowering workflow-based network analysis. *Genome Biol.* **20**, 1–15, 2019). Additional node and edge attributes, such as gene functional annotations and interaction confidence scores, were added for a comprehensive view. For layout optimization and clarity, Perfuse Force-Directed layout was applied in Cytoscape 3.10. Layout parameters were fine-tuned to reflect interaction strengths, with edge weights influencing node proximity, which effectively clusters key regulators and functional modules. Adjustments to repulsion and gravity settings minimized node overlap, maintaining cohesion across the network. To emphasize prominent nodes, color-coded nodes were added based on functional annotations and node size scaled based on degree centrality, highlighting central regulatory hubs. Edge thickness was similarly adjusted to indicate interaction confidence, with thicker edges representing stronger regulatory links. Community detection and centrality analyses within Cytoscape 3.10 were used to pinpoint top regulators and functional modules, which supported a more detailed interpretation of the network’s structural and functional organization.

### Genetic variation and geoclimatic variable associations

Association between genetic variation of bHLH35 and geoclimatic variables were downloaded from the Arabidopsis GenoClim 2.0 database^19^. The scores for association strength (negative logarithm of the P-value) computed by a mixed model that corrects for population structure (AMM_scores) were plotted in the heatmap with the indicated q-values corrected by the Benjamini-Hochberg method.

### Weighted Gene Co-expression Network Analysis

To identify and analyze co-expressed genes, we performed Weighted Gene Co-expression Network Analysis (WGCNA) in R^30^. Optimal soft-thresholding power was determined to be 18 by the pickSoftThreshold function in WGCNA giving a scale-free topology fit index (SFT.R²) > 0.8 and a mean connectivity (mean.k) <100. The weighted co-expression network was then constructed by the blockwiseModules function. The input matrix consisted of the 47 replicates and 9,260 genes, with expression values transformed using variance-stabilizing transformation (VST) from DESeq2 to normalize the count data. The parameters for blockwiseModules were: 1) soft threshold power of 18, as established by pickSoftThreshold; 2) “signed” network and Topological Overlap Matrix (TOM) types to preserve correlation directionality; 3) minimum module size of 40 genes; 4) module merging threshold of 0.18 (to merge highly similar modules). This resulted in a WGCNA network comprising 21 modules (Table S11).

### Transcription factor (TF) binding site enrichment within WGCNA co-expression modules

To analyze the promoter sequences of the genes in our WGCNA network, we utilized the TAIR10 genome assembly of *Arabidopsis thaliana* along with gene annotations from Ensembl Plants release 49 (https://plants.ensembl.org/Arabidopsis_thaliana/Info/Index). Promoter regions were defined as spanning 1 kb upstream to 300 bp downstream of the transcription start site (TSS). Validated TF binding profiles were retrieved from JASPAR2024 (Rauluseviciute, I. JASPAR 2024: 20th anniversary of the open-access database of TF binding profiles. *Nucleic Acids Res.* **52**, D174–D182, 2024). Connection to JASPAR2024 was made using the RSQLite package (Müller, K., Wickham, H., James, D. A. & Falcon, S. RSQLite: SQLite Interface for R. R package version 2.3.9, 2024 https://github.com/r-dbi/RSQLite, https://rsqlite.r-dbi.org.) and Position Weight Matrices (PWMs) corresponding to the *Arabidopsis thaliana* species code (3702) were retrieved by TFBSTools (Tan, G. & Lenhard, B., TFBSTools: an R/bioconductor package for TF binding site analysis. *Bioinformatics* **32**, 1555–1556, 2016). These PWMs served as the basis for downstream motif enrichment analysis. Motif enrichment within WGCNA gene modules was computed using the calcBinnedMotifEnrR function from the monaLisa R package (Machlab, L. et al. monaLisa: an R/Bioconductor package for identifying regulatory motifs. *Bioinformatics* **38**, 2624–2625, 2022). To ensure that motif enrichment was evaluated within biologically relevant co-expression networks, we assigned each promoter sequence to its respective WGCNA co-expression module, creating module-specific bins. Enrichment calculations were performed relative to all promoter sequences across all modules by setting the background to “allBins.” This approach allowed us to assess the overrepresentation of TF binding motifs within each module’s promoters while accounting for background motif frequencies, thereby identifying WGCNA modules potentially enriched for specific TF targets. This binned motif enrichment analysis returned the FDR-adjusted p values (-log10 scale) of the enrichment of each TF motif in each bin (WGCNA module) relative to its occurrences in the background sequences.

### Hierarchical and co-regulating models

TF regulators for the modules of interest (blue, magenta, tan) were identified as TFs whose motifs were enriched for the genes in the module at adjusted p value threshold of 0.05. To find the matches of specific TF motifs in target gene promoters, we used the findMotifHits function in monaLisa with the query parameter set to the position weight matrices (PWMs) of the TFs of interest, the subject parameter defined as the set of promoter sequences corresponding to all genes within the WGCNA module of interest, and motif matching method set to the matchPWM method at a minimum score threshold of 6.0. The visualizations were done with Cytoscape (Shannon, P. Cytoscape: a software environment for integrated models of biomolecular interaction networks. *Genome Res.* **13**, 2498–2504, 2003).

### Expression heatmaps and GO analysis and heatmaps

The expression heatmaps were created with the ComplexHeatmap package in R (Gu, Z. Complex heatmap visualization. Imeta 1, e43, 2022). Every gene was centered and scaled along all 47 samples, so gene expression is represented as the z-score of the VST-normalized count data. K-means clustering of the samples and genes were performed by setting the row_km and column_km parameters in the Heatmap function. The Gene Ontology (GO) enrichment analysis utilized the compareCluster function from the clusterProfiler R package (Yu, G., Wang, L. G., Han, Y. & He, Q. Y. clusterProfiler: an R package for comparing biological themes among gene clusters. *OMICS* **16**, 284–287, 2012). These dotplots identified the Biological Process (BP) terms that were significantly associated with our modules of interest.

### Whole-genome transcription factor yeast two-hybrid assay

To identify TF partners of bHLH35, we implemented a high-throughput yeast two-hybrid (Y2H) screen using Arabidopsis bHLH35 as bait against a whole-genome Arabidopsis TF library described by^25^. Coding sequence (CDS) of bHLH35 was cloned into a modified pGBKT7 vector harboring the GAL4 DNA binding domain (DNA-BD) where the URA marker was replaced by a HIS selection marker using In-Fusion cloning kit (Takara, CA, USA; 638955). The resulting DNA-BD-bHLH35 plasmid was transformed into the Yeast 2 Gold (Y2G) yeast strain (Takara, CA, USA; 630498) using the lithium acetate method^46^. Y2G cells carrying the CDS of bHLH35 were mated with the Yα1867 strain carrying the Arabidopsis TFs library and diploid strains were selected on YNB-His-Ura-Trp+Ade media (Sunrise Science Products, TN, USA; 1500, 1051). This Y2H screen relies on the Ade marker to detect protein-protein interactions; therefore, after selecting positive mating colonies carrying both the bait and preys, diploid strains were pinned on quadruple drop out media (YNB-His-Ura-Trp-Ade; Sunrise Science Products, TN, USA; 1500, 1159) using a HDA RoToR robot (Singer Instruments, UK) and after two days of incubation, white colonies were selected for further confirmation through a secondary screen and Sanger sequencing. Details of the primers used for cloning are listed in Table S10.

### Yeast one-hybrid (Y1H) assay

To test whether bHLH35 binds to the promoter region of *LBD31* (AT4G00210), we performed a targeted yeast one-hybrid (Y1H) assay. Briefly, the CDS of bHLH35 was cloned into a pENTR vector using In-Fusion cloning kit (New England Biolabs, Ipswich, MA, USA). The sequence-verified pENTR-bHLH35 construct was shuttled into the destination vector pDEST22 by Gateway LR recombination reaction (Invitrogen, CA, USA; 11791020). Next, promoter regions comprising 500 bp upstream of the transcription start site and 183 bp 5’ UTR were cloned as two fragments called LBD31-Promoter 1 (183 bp 5’ UTR + 200 bp promoter upstream of the transcription start site) and LBD31-Promoter 2 (300 bp promoter; 200 bp to 500 bp upstream of the transcription start site). The two fragments (Table S10) were amplified from *A. thaliana* genomic DNA using a high-fidelity polymerase (NEB, MA, USA; MO530S) and cloned into the pHIS plasmid using the In-Fusion cloning kit following the manufacturer’s protocols. The pHIS plasmid containing the promoter fragments was linearized with XhoI and integrated into the yeast Y1H-aS2 (his3-Δ1) kindly provided by the Walhout lab as described by^47^. Positive colonies were selected on minimal media deficient in His (YNB-His, Sunrise Science Products, TN, USA; 1006). Then, the Y1H-aS2 strains carrying the *LBD31* promoter fragments were transformed with pDEST22-*bHLH35*, and positive colonies were selected on YNB-His-Trp media (Sunrise Science Products, TN, USA). The interaction of bHLH35 with *LBD31* promoter fragments was tested on selective media at varying concentrations of 3-aminotriazole (3-AT, Thermo Fisher Scientific, MA, USA; 264571000). Details of the primers used for cloning and the promoter fragment sequences are listed in Table S10.

### Protein isolation

Coding sequences of the *bHLH35* (AT5G57150) and *NAC069* (AT4G01550) were cloned into pET28a+ bacterial gene expression vectors downstream of the 6xHIS tag. The clones and empty vector were transformed along with a vector containing molecular chaperones pG-KJE8 (Takara, CA, USA; 3340) into *E. coli* BL21 strain and transformants were selected on LB plates containing Kanamycin (50 µM/mL, Gold Bio, MO, USA; K-120-5) and Chloramphenicol (25 µM/mL, Gold Bio, MO, USA; C-105-5). Single colonies were inoculated in 3 mL LB containing Kanamycin (50 µM/mL) and Chloramphenicol (25 µM/mL) and grown overnight at 37 °C at 180 rpm. 50 mL LB was inoculated with 500 µL of primary overnight culture supplemented with Tetracycline (8 ng/mL, Gold Bio, MO, USA; T-101-25), L-Arabinose (2.5 mg/mL; Sigma-Aldrich, MO, USA; A-3256), Kanamycin (50 µM/mL) and Chloramphenicol (25 µM/mL). Cultures were incubated at 37 °C at 180 rpm until an OD_600_ of 0.3. To a final volume of 50 mL culture (0.3 OD_600_), 0.5 mM IPTG (Gold Bio, MO, USA; I2481C) was added and incubated at 16 °C at 150 rpm for 20 hours. The bacterial culture after 20 hours of incubation was centrifuged at 6000 rpm for 10 minutes, cell pellets were resuspended in 4 mL lysis buffer (50 mM Tris-HCl pH 7.5, 50 mM NaCl, 10 mM Imidazole, broad-spectrum Pierce™ Protease Inhibitor, Thermo Fisher Scientific, MA, USA; A32955) and sonicated 20 times for 20 seconds, with 20 seconds interval in between (Ultrasonic processor 130 watt, 30 kHZ, cv18 converter probe), in ice. Lysates were centrifuged at 12000 rpm for 20 minutes, supernatant was transferred to 15 mL falcon tube and incubated with 50 µL of HIS Ni-NTA Agarose resin (Qiagen, MD, USA; 30210), pre-calibrated with wash buffer (50 mM Tris-HCl pH 7.5, 50 mM NaCl, 50 mM Imidazole, broad-spectrum Pierce™ Protease Inhibitor), at 4 °C for 2 hours on tube rotator. The protein extracts and resin mix were spun down at 500 rpm for 30 seconds. The supernatant (protein soup) was carefully removed, and resins were incubated with ice-cold wash buffer (50 mM Tris-HCl pH 7.5, 50 mM NaCl, 50 mM Imidazole, Pierce™ Protease Inhibitor) at 4 °C for 5 minutes (shaking). The washing step was repeated three times with 5 mL wash buffer every time. Protein was eluted with 200 µL elution buffer (50 mM Tris-HCl pH 7.5, 50 mM NaCl, 250 mM Imidazole, Protease Inhibitor). Eluted protein was mixed with glycerol (10% final concentration) and stored at −80 °C until further use. Protein quantification was performed using Bradford reagent (Thermo Fisher Scientific, MA, USA; 23200). Details of the primers used for cloning are in Table S10.

### Electro-Mobility Shift Assay (EMSA)

EMSA analysis was performed with biotin-labeled LBD31 promoter probes and LBD31 mutant promoter probes (E-box replaced with random sequences; Table S10) using LightShift® Chemiluminescent EMSA Kit (Thermo Fisher Scientific, MA, USA; 20148) following the manufacturer’s protocol. For the binding reaction, probes (Table S10) were incubated with bHLH35 and/or NAC069 protein (isolated as described above) in 1X binding buffer and 2.5% glycerol. Probe-protein mix was separated on 6% Polyacrylamide gel, transferred to nylon membrane (Amersham Hybond^TM^-N, GE Healthcare, IL, USA) and cross-linked at 120 mJ/cm^2^ (UV-Crosslinker, FB-UVXL-1000, Fisher Scientific, NH, USA). Chemiluminescence-based detection of biotin-streptavidin conjugate was performed following the LightShift® Chemiluminescent EMSA Kit’s protocol instructions. Details of probes are in Table S10.

### Mutant Complementation

The *bhlh35_2* mutant was transformed with the *bhlh35* CDS driven by the native *bHLH35* promoter (1000 base fragment). The promoter of the *bHLH35* gene (AT5G57150), and coding sequences of mGFP and *bHLH35* (in the following order: *bHLH35* promoter-mGFP-bHLH35 gene) were cloned into pCAMBIA1302 binary vector. Cloning was performed using in-fusion cloning kit (Takara, CA, USA; 638955). After confirming the sequence, the recombinant vector was transferred to *Agrobacterium tumefaciens* GV3101. *Arabidopsis* plants were transformed by floral dip. T1 seeds were screened on ½ MS media supplemented with 12 μg/mL hygromycin as described in^48^. Ten different homozygous transgenic lines were generated (T3) and tested for survival under conditions of stress combination as described above.

### Data analysis

Two-tailed Student’s t-test was used to determine significant differences between the percent survival of WT and the different mutant genotypes. The mean percent survival difference at a p-value ≤ 0.05 was considered as significant.

## Supplementary Material

### Supplementary Figure Legends

**Fig. S1.**
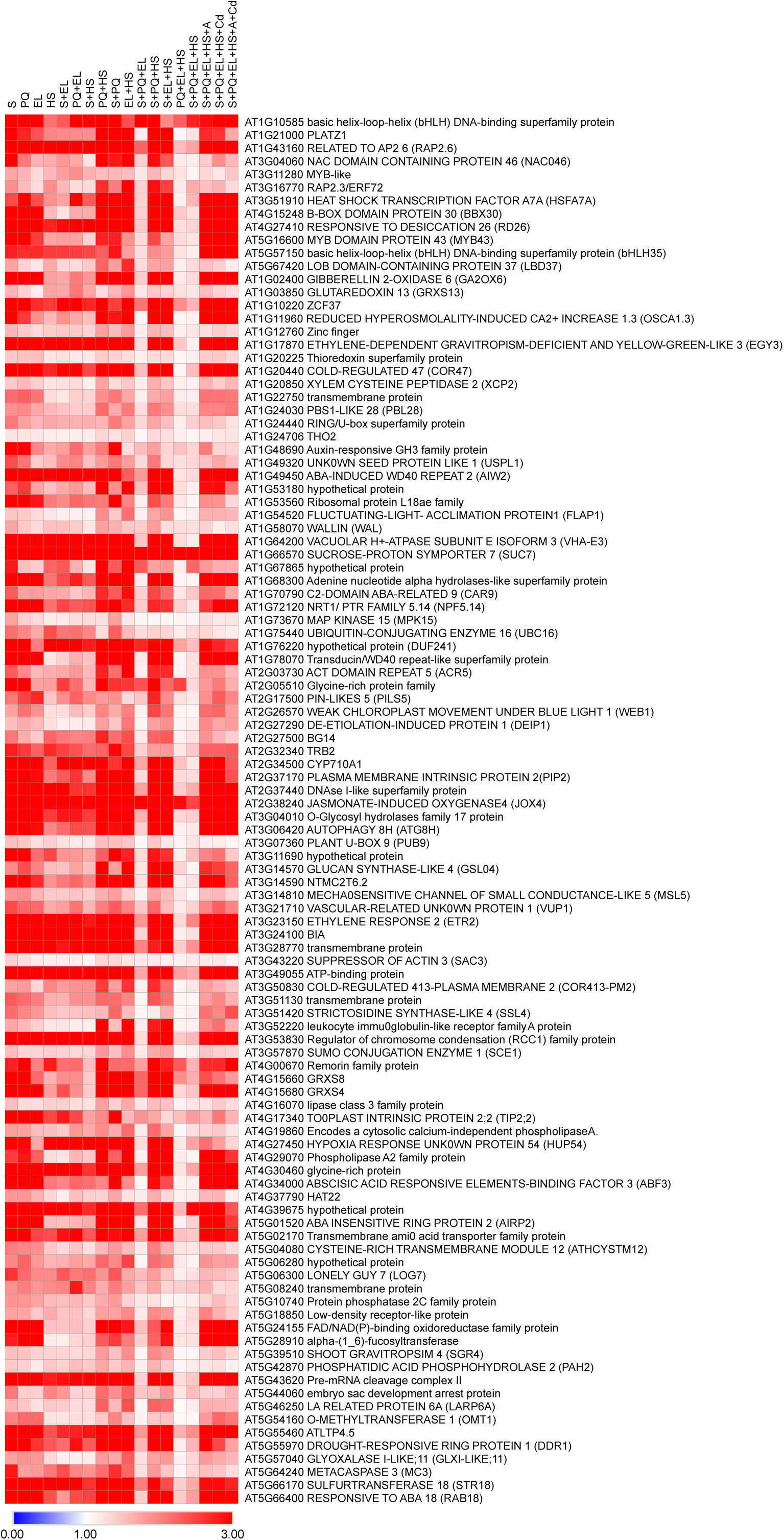
Expression of transcripts significantly elevated in response to a combination of 2-, 3-, 4-, 5-, and 6-different abiotic stresses. Heatmap showing the expression of all 130 transcripts with significantly enhanced expression under conditions of multifactorial stress combination (Salt, S; the herbicide paraquat, PQ; cadmium, Cd; excess light, EL; and heat stress, HS; Acidity, A). In support of Fig. 1.

**Fig. S2.**
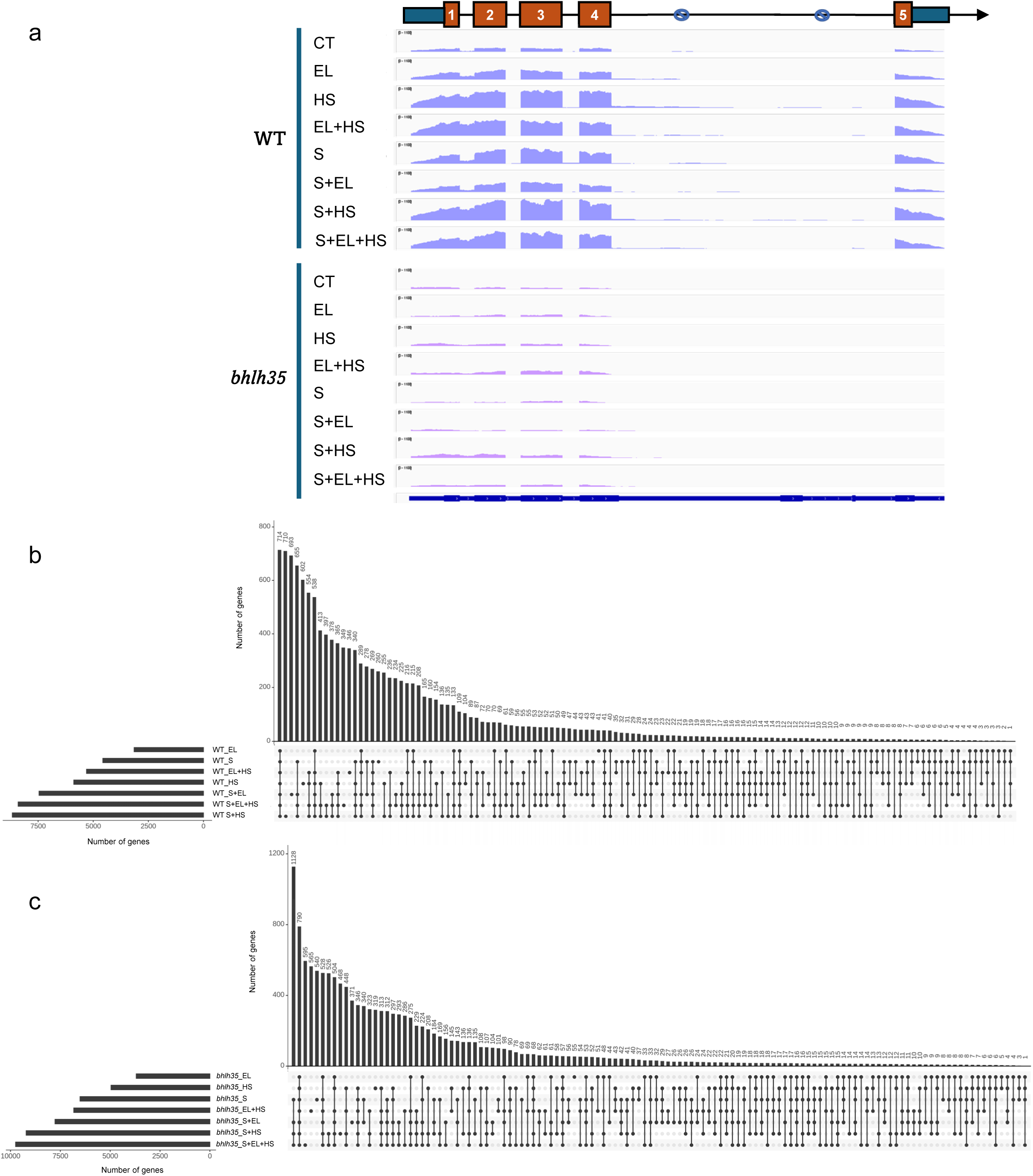
RNA-Seq analysis of wild type (WT) and the *bhlh35_2* mutant subjected to control (CT), and Salt (S), excess light (EL) and heat stress (HS), in all possible combinations. **a,** RNA-Seq read maps aligned to the bHLH35 gene from plants subjected to all stress treatments. Note that intron 1 is not spliced under conditions of HS. **b,** and **c**, Upset plot showing the overlap between transcripts significantly altered in their expression in wild type (WT; b), or the *bhlh35_2* mutant (c) in response to CT and all stress treatments. In support of Figs. 1 and 2.

**Fig. S3.**
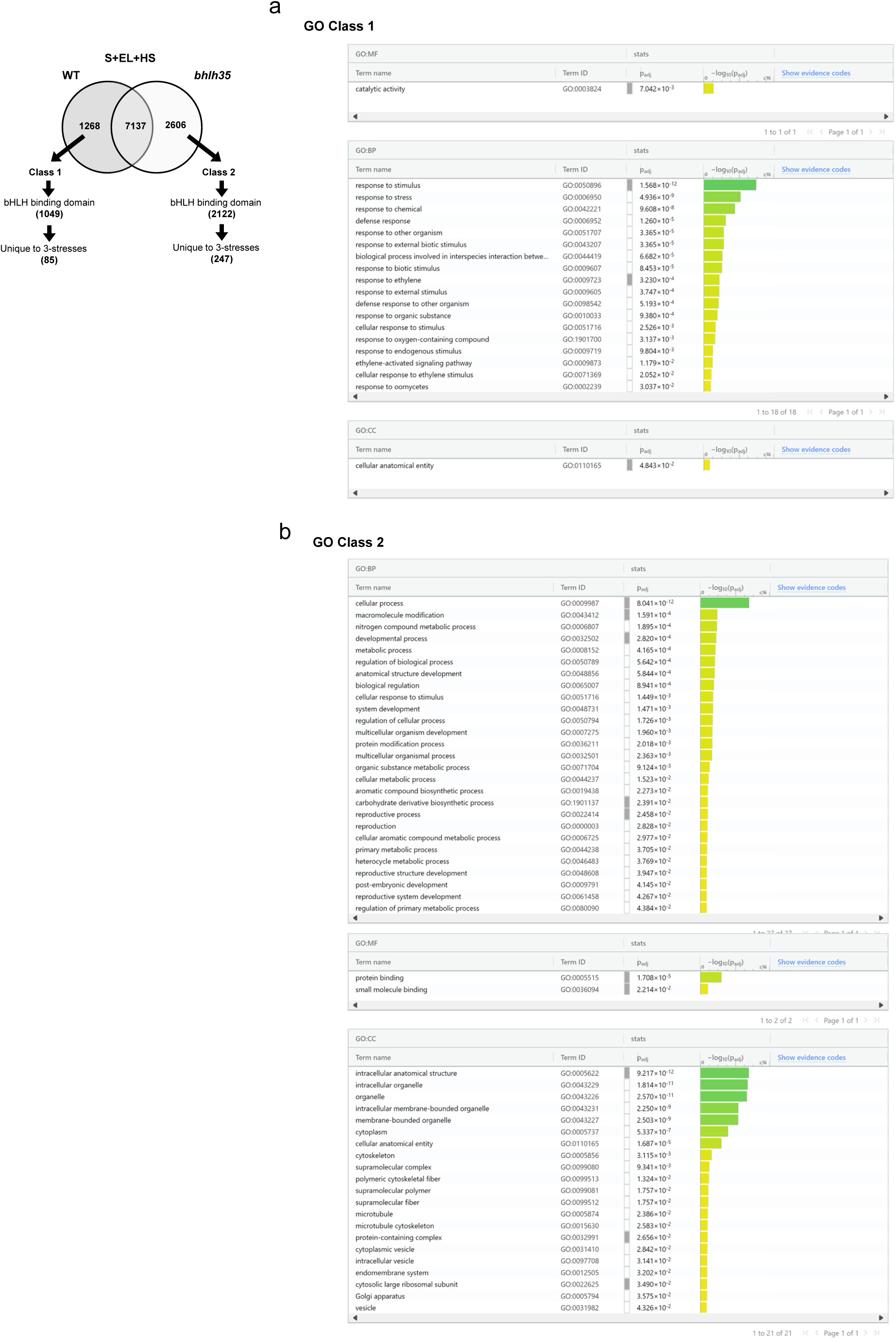
Gene ontology (GO) annotation for transcripts significantly altered in their expression in wild type (WT) and the *bhlh35_2* mutant in response to a combination of S+EL+HS (Class 1 and Class 2). **a,** Venn diagram from Fig. 2 showing the two classes. **b**, GO annotations of the two classes. In support of Fig. 2.

**Fig. S4.**
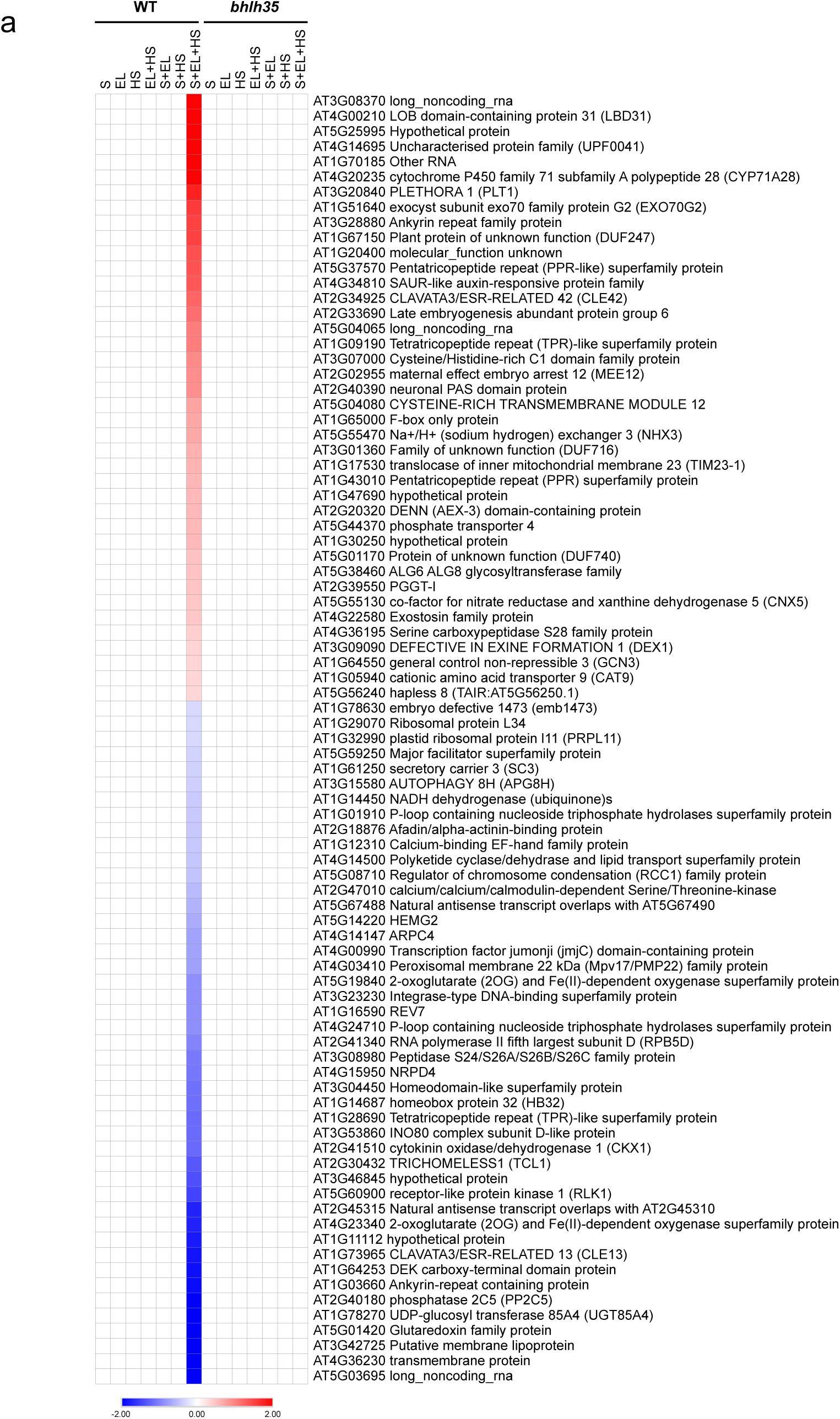
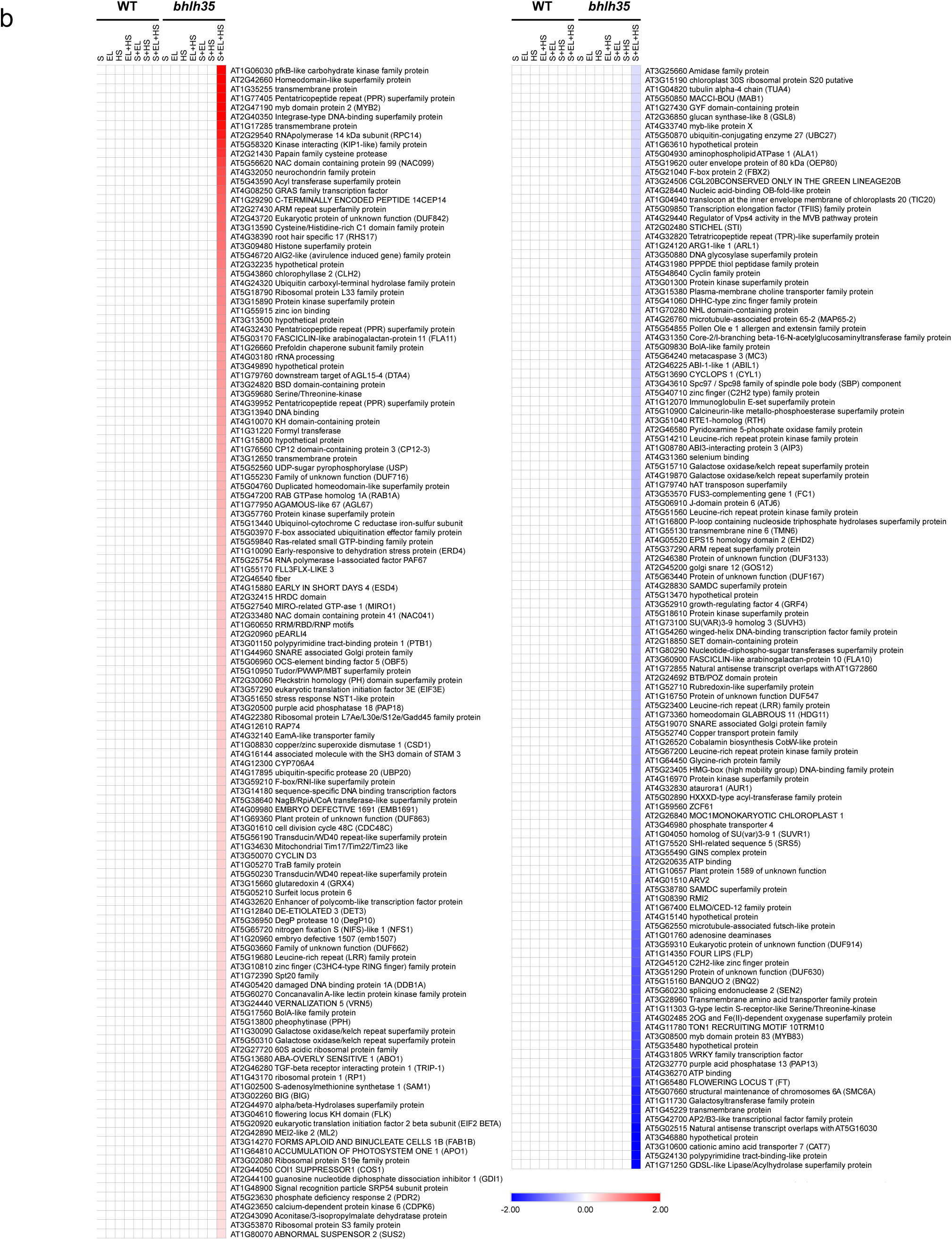
Heatmaps showing the expression of transcripts with a pattern unique to wild type (WT) or the *bhlh35_2* mutant in response to Salt (S), excess light (EL) and heat stress (HS), in all possible combinations. **a,** Expression of transcripts unique to a combination of S+EL+HS in WT but not the *bhlh35_2* mutant. **b**, Expression of transcripts unique to a combination of S+EL+HS in the *bhlh35_2* mutant but not WT. In support of Fig. 2.

**Fig. S5.**
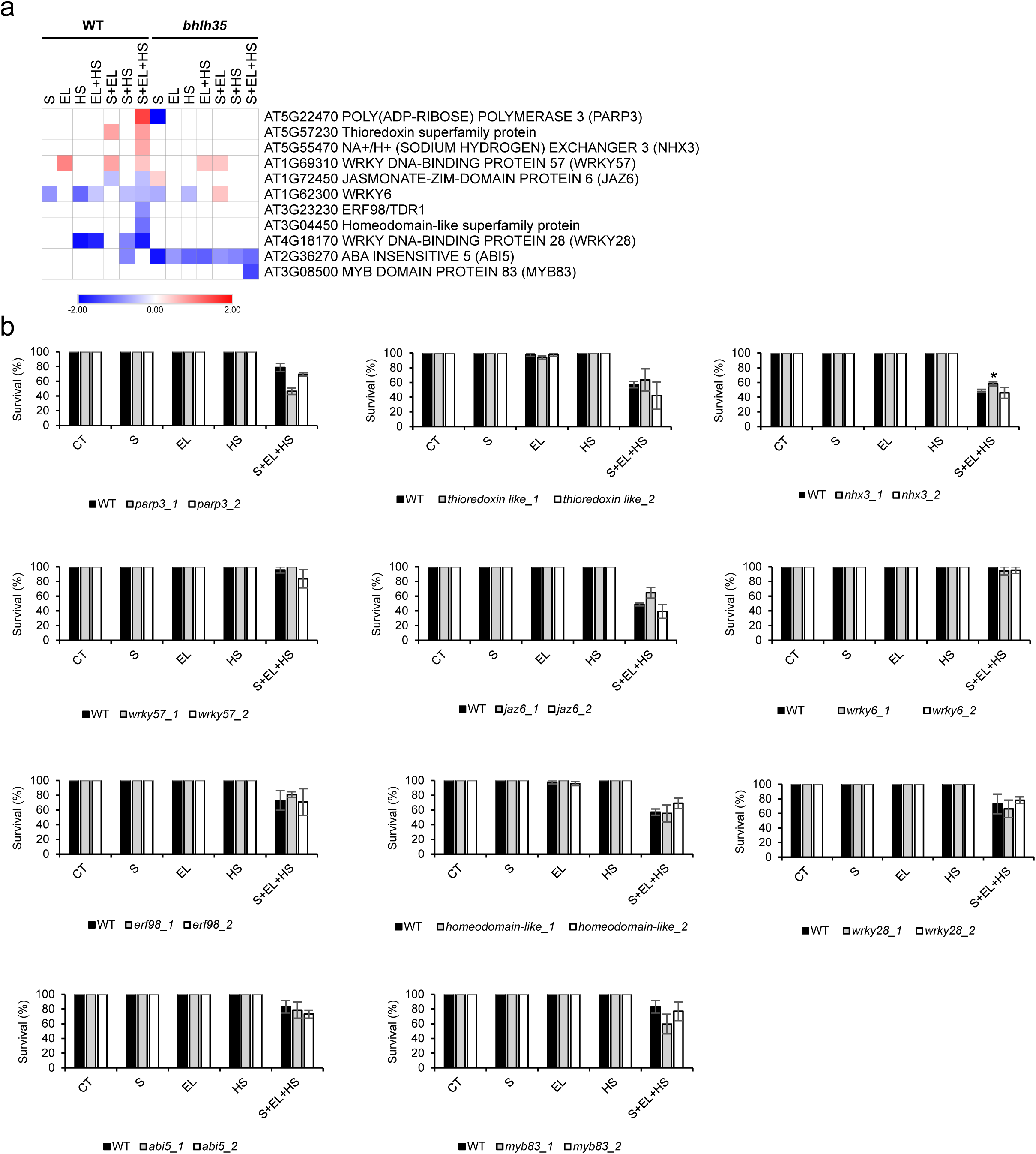
Survival of two independent mutants for different genes randomly selected from Class 1 and Class 2 transcripts (Fig. 2a) having *bHLH35-*dependent expression under conditions of Salt (S) + excess light (EL) + heat stress (HS) (S+EL+HS). **a,** Heatmap showing expression of transcripts that depend on *bHLH35* under a combination of S+EL+HS. **b**, Survival of two independent mutants for each of the genes encoding transcripts that require *bHLH35* for expression under conditions of S+EL+HS (From a). *Two-tailed Student’s t-test (p ≤ 0.05). In support of Fig. 3.

**Fig. S6.**
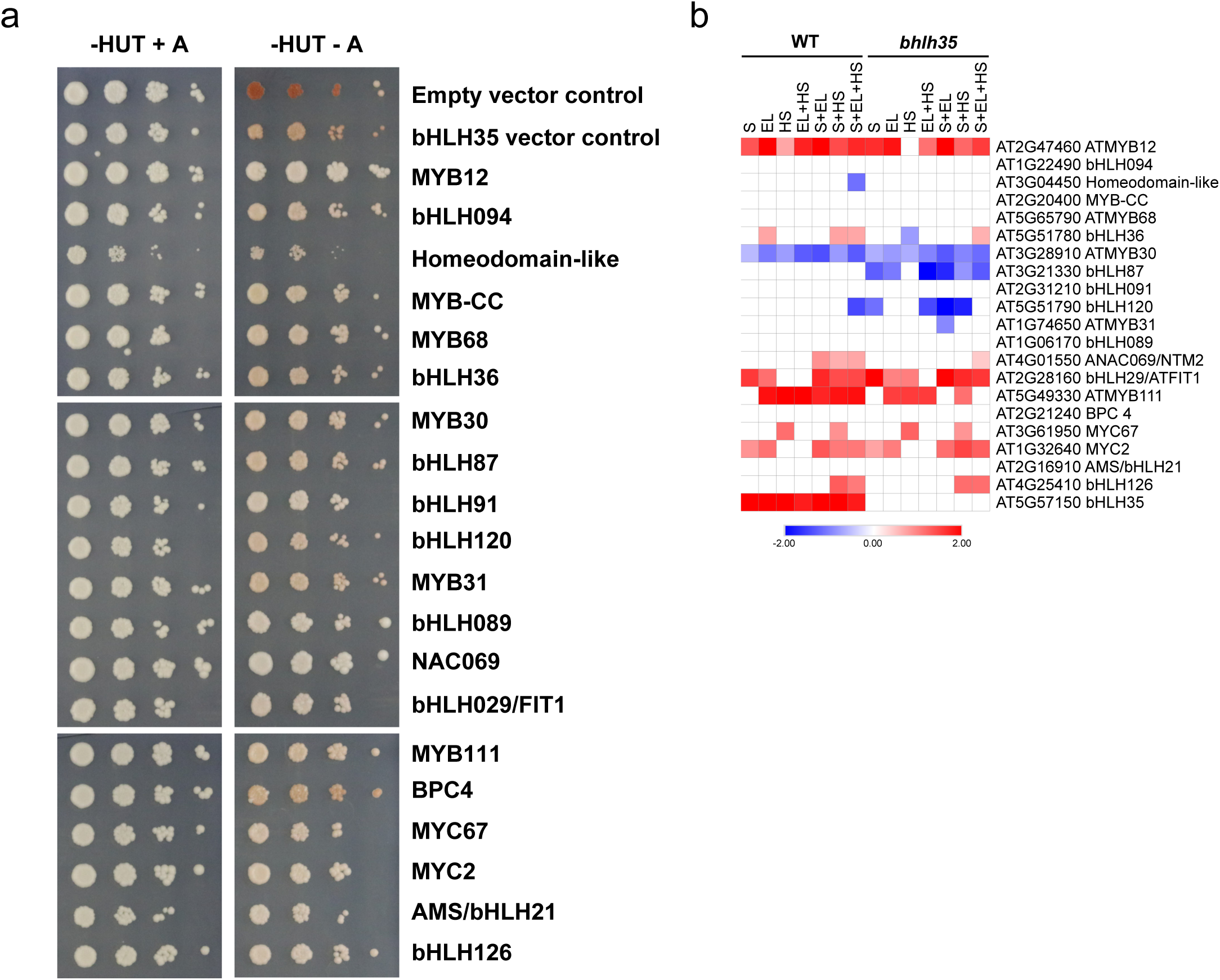
Yeast 2-hybrid analysis of transcription factors (TFs) that interact with bHLH35. **a**, Transcription factors (TFs) found to interact with bHLH35. **b**, Heatmap showing the expression of transcripts encoding the TFs identified in (a) under conditions of salt (S), excess light (EL), and heat stress (HS) in all possible combinations. In support of Fig. 3.

**Fig. S7.**
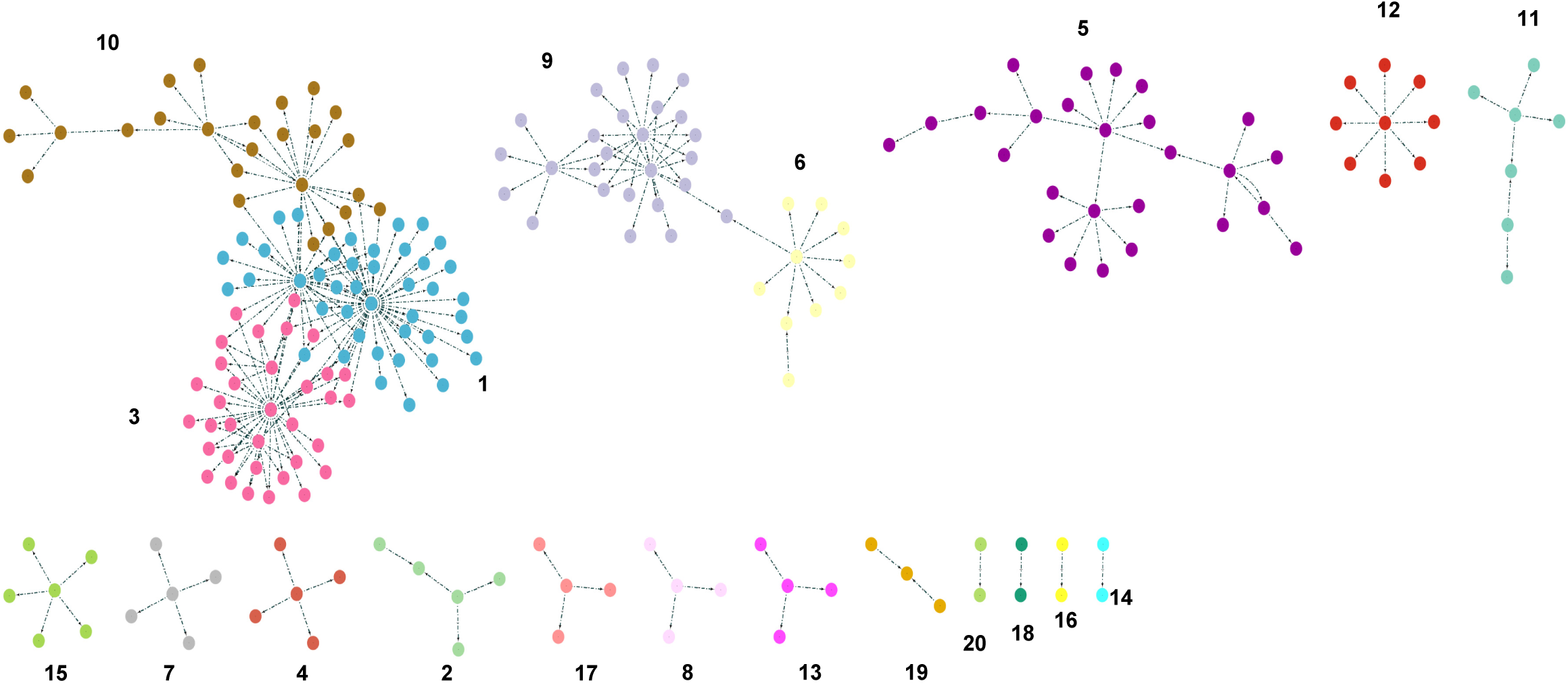
Gene regulatory network (GRN) analysis of *bHLH35.* Gene regulatory network analysis (using Differential Inference Analysis of Expression; DIANE) of *bHLH35* based on the RNA-Seq results obtained in this study and other available resources and databases. Identified pathways and description of each node associated with *bHLH35* are in Table S9. In support of Fig. 4.

**Fig. S8a.**
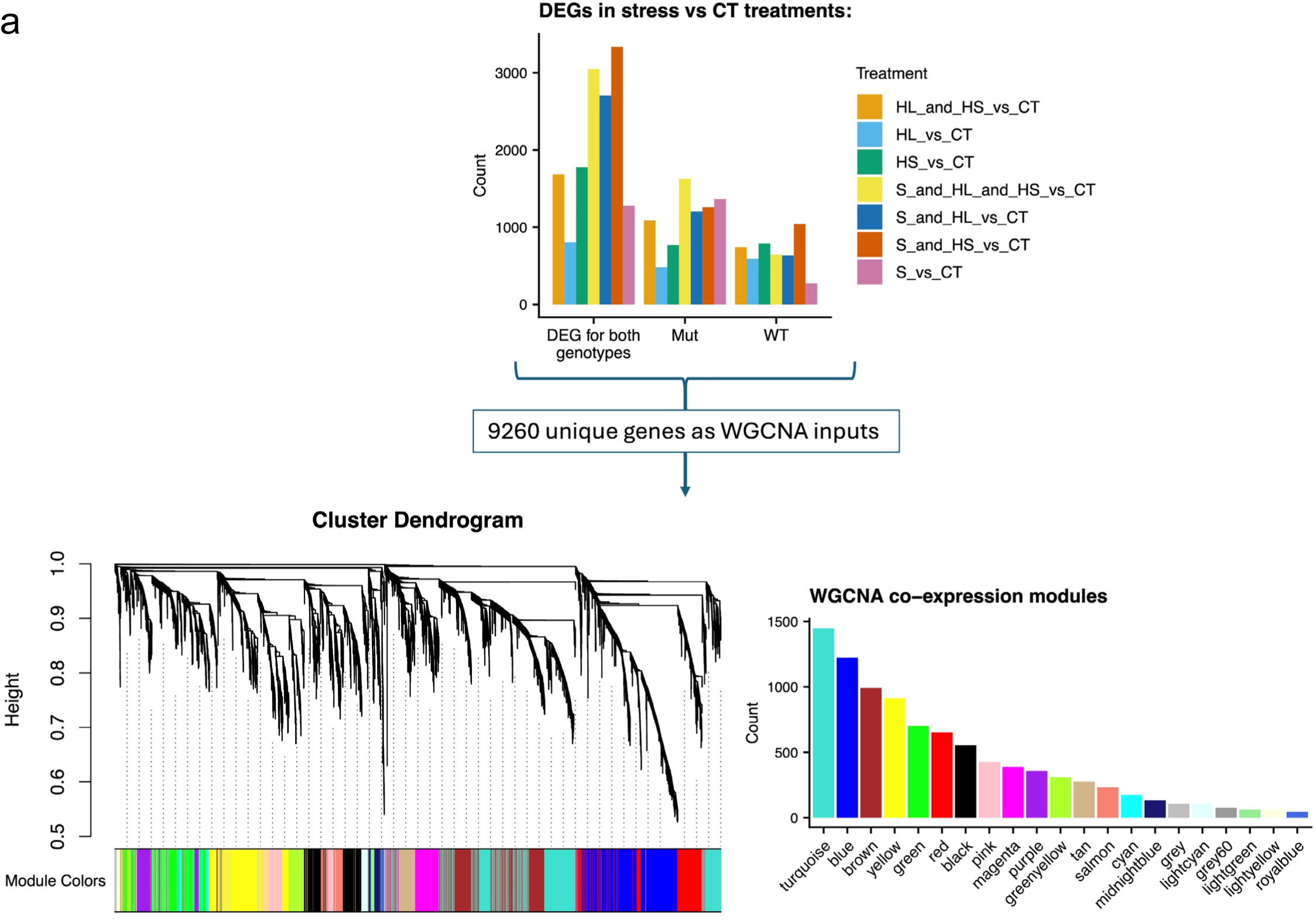
9260 unique genes were identified as DEGs and their expression used as input to create a WGCNA network of 21 co-expression modules. For each of the seven stress conditions, genes were considered differentially expressed if the control versus stress condition had a fold change greater than 1.8 and an FDR-adjusted p value less than 0.01. Thus, we took the union of the DEGs from the WT samples in CT versus stress conditions and the DEGs from the *bhlh35* mutant samples in CT versus stress conditions, resulting in a total of 9260 unique genes. The expression of these 9260 genes for the 47 samples was used as input for WGCNA that created a co-expression network of 21 modules marked by different colors. In support of Fig. 4d.

**Fig. S8b.**
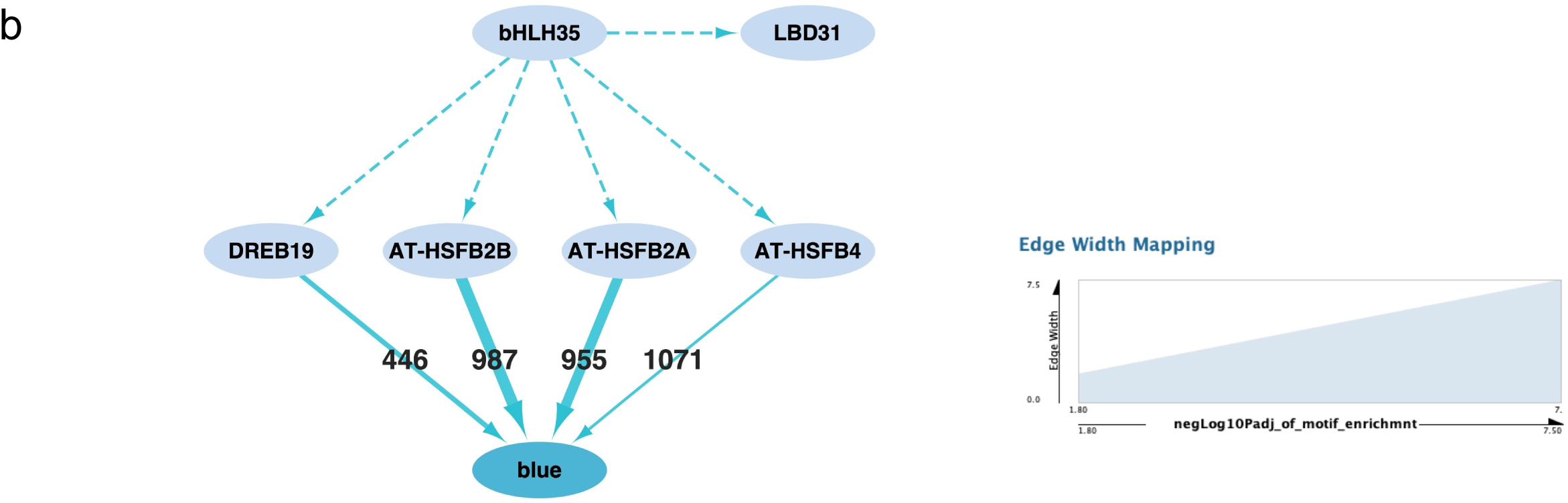
Model of hierarchical transcriptional regulatory network of genes in the blue module regulated by bHLH35. bHLH35 and LBD31 are co-expressed in the blue module with DREB19, AT-HSF2B, AT-HSFB2A and AT-HSFB4. The promoter sequences of LBD31, DREB19, AT-HSF2B, AT-HSFB2A and AT-HSFB4 contain multiple matches of binding site motifs from the bHLH TF family (dashed arrows). The binding site motifs for DREB19, AT-HSF2B, AT-HSFB2A and AT-HSFB4 were significantly enriched in the promoter sequences of the blue module genes, as indicated by solid arrows labeled by the number of genes that contained the binding site motifs and the thickness scaled to the -log10 enrichment p-values corrected by FDR. In support of Fig. 4d.

**Fig. S8c.**
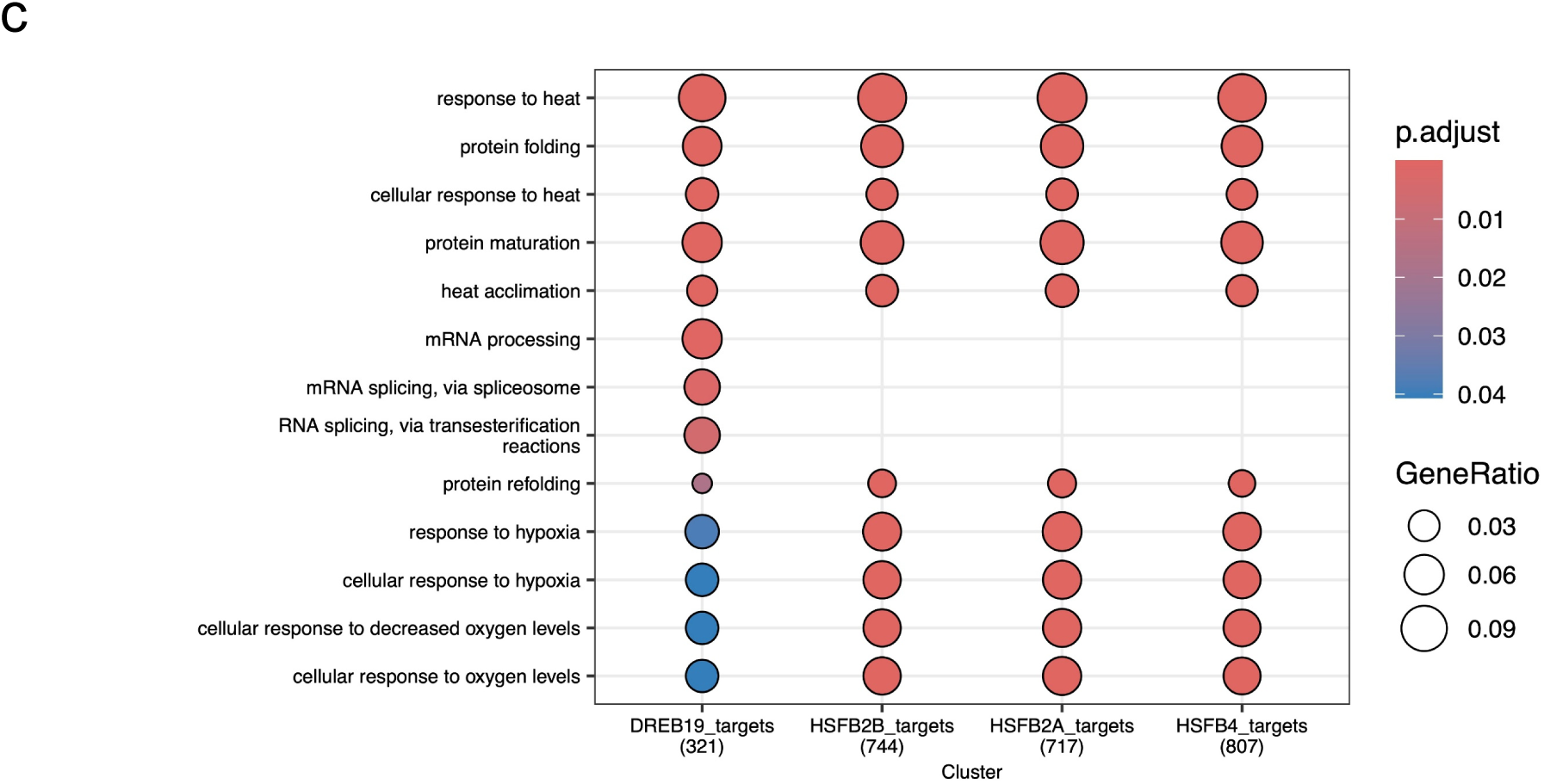
The four TFs with motifs significantly enriched in the blue module, and co-expressed with bHLH35, seem to target genes associated with heat and hypoxia responses. The genes in the blue module with the corresponding TF motifs in their promoter region (thus potential TF target genes) were significantly enriched for biological processes related to heat, protein folding, and oxygen/hypoxia events. Most of these GO terms are shared by the potential targets of these TFs. In support of Fig. 4d.

**Fig. S8d.**
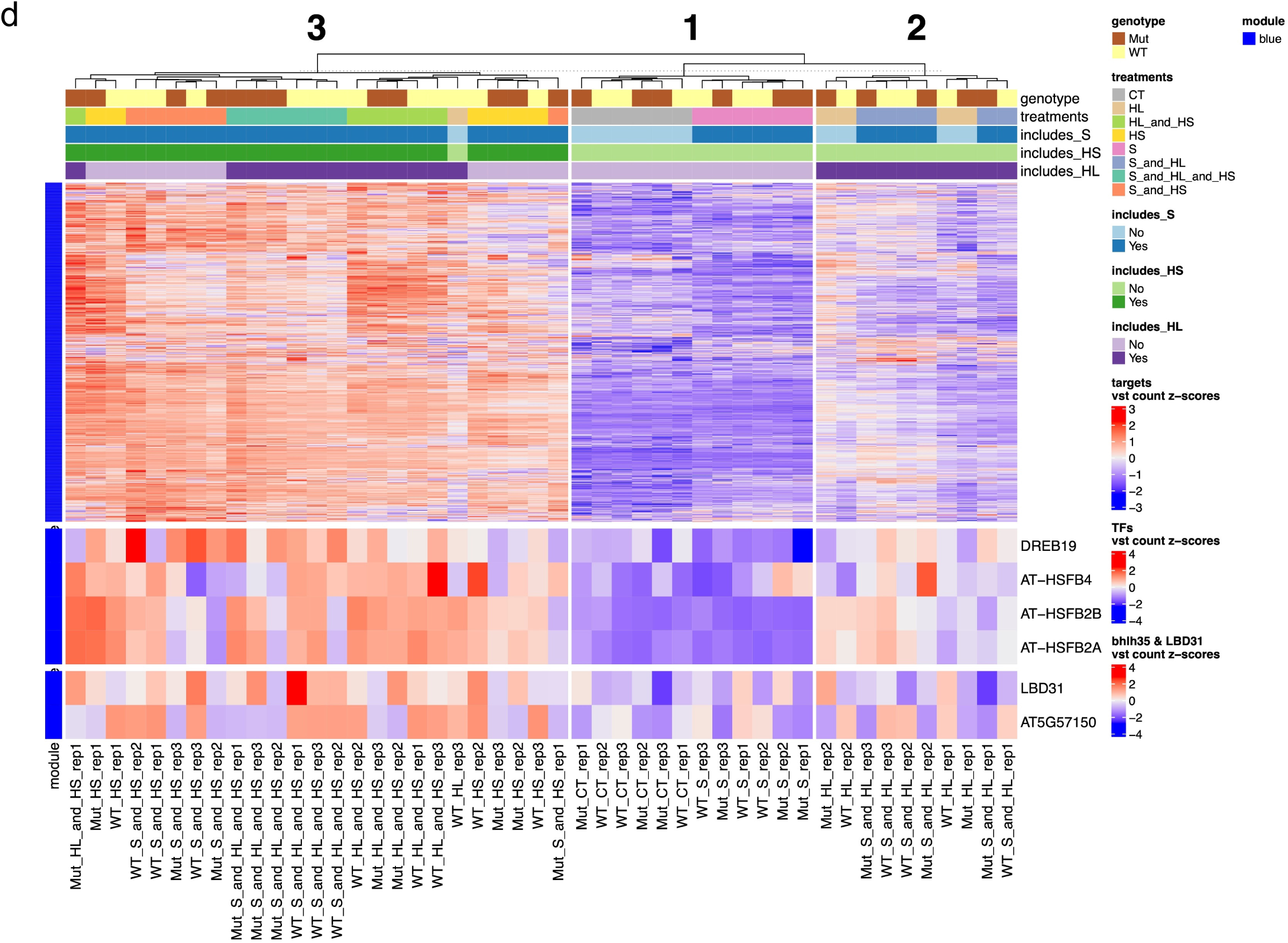
LBD31, bHLH35 (AT5G57150), the four enriched TFs (DREB19, AT-HSFB4, AT-HSFB2A, AT-HSFB2B) and their target genes demonstrate correlated expression patterns – validating their membership in the same co-expression module (blue module). As seen in the expression heatmap, LBD31, bHLH35, the 4 TFs and their targets all seemed to follow a similar pattern of downregulation for all the treatments that did not include heat stress (HS). Specifically, the genes were least expressed (most negative z-scores) in the control condition (CT) and the salt condition (S) regardless of genotype for the WT and mutant samples (cluster 1). Cluster 2 contains both mutant and WT samples from the high light (HL/EL) as well as the ‘S and HL/EL’ conditions. The genes in these two treatments were for the most part downregulated but to a lesser extent than the ‘CT’ and ‘S’ cluster. Finally, the third and largest cluster (cluster 3) exhibited a pattern of upregulation (most positive z-scores) and grouped together all the treatments that included HS (except for one outlier, WT_HL_rep3). This is coherent with our proposed model as it contains the three heat stress TFs that target a large majority of the genes in the blue module, thereby explaining why the treatments that included HS led to major upregulation of these genes. In support of Fig. 4d. Acronyms: HS = heat stress; HL = high light; S = salt.

**Fig. S8e.**
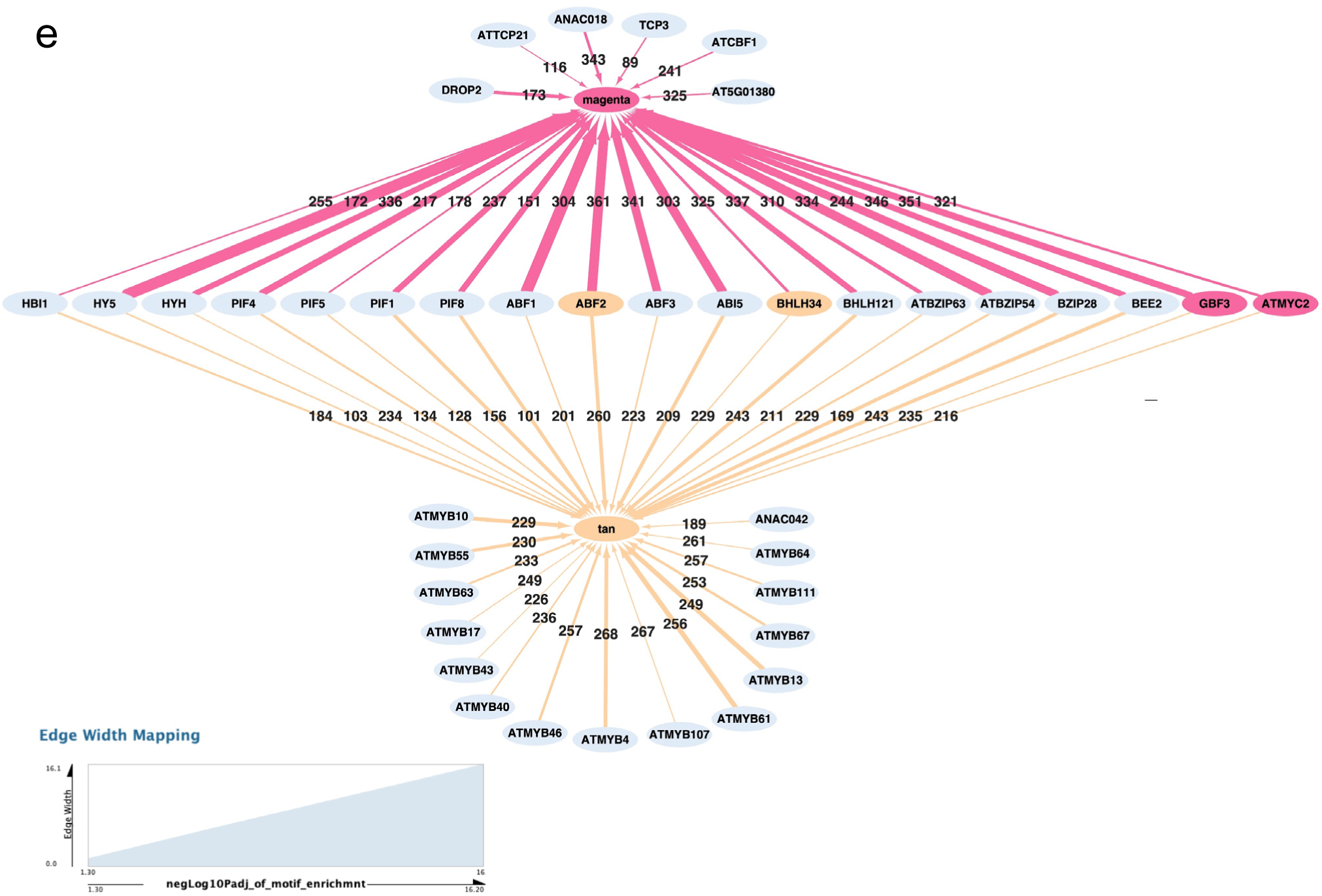
Model of bHLH35 coregulated modules. Although the JASPAR 2024 database does not contain a motif bHLH35, we found significant enrichment for the nine Arabidopsis bHLH TF motifs that are in the database (bHLH3, bHLH13, bHLH121, bHLH34, bHLH74, bHLH18, bHLH77, bHLH72 and bHLH49) in the promoter sequences of genes in the tan and/or magenta modules (FDR-adjusted p <0.05), raising the possibility that bHLH35 could be targeting a significant proportion of genes in these two modules. Enrichment analysis using all the Arabidopsis TFs in the JASPAR2024 database identified 40 TFs that are present in the WGCNA network and had enriched motifs in the promoters of the genes in these two modules as potential co-regulators. Of note, although these 40 TFs were themselves present in the WGCNA network, they belonged to different modules– except ABF2 and bHLH34 which belonged to the tan module, and GBF3 and ATMYC2 which belonged to the magenta module. We further filtered all the target genes in the tan and magenta modules and removed those that did not have bHLH binding motifs within their promoters for GO enrichment analysis (panel f) and expression clustering (panel g). The numbers on the edges indicate the number of target genes and the thickness of the edge is scaled by the -log10 FDR adjusted p-values of the motif enrichment for the genes in the module. In support of Fig. 4d.

**Fig. S8f.**
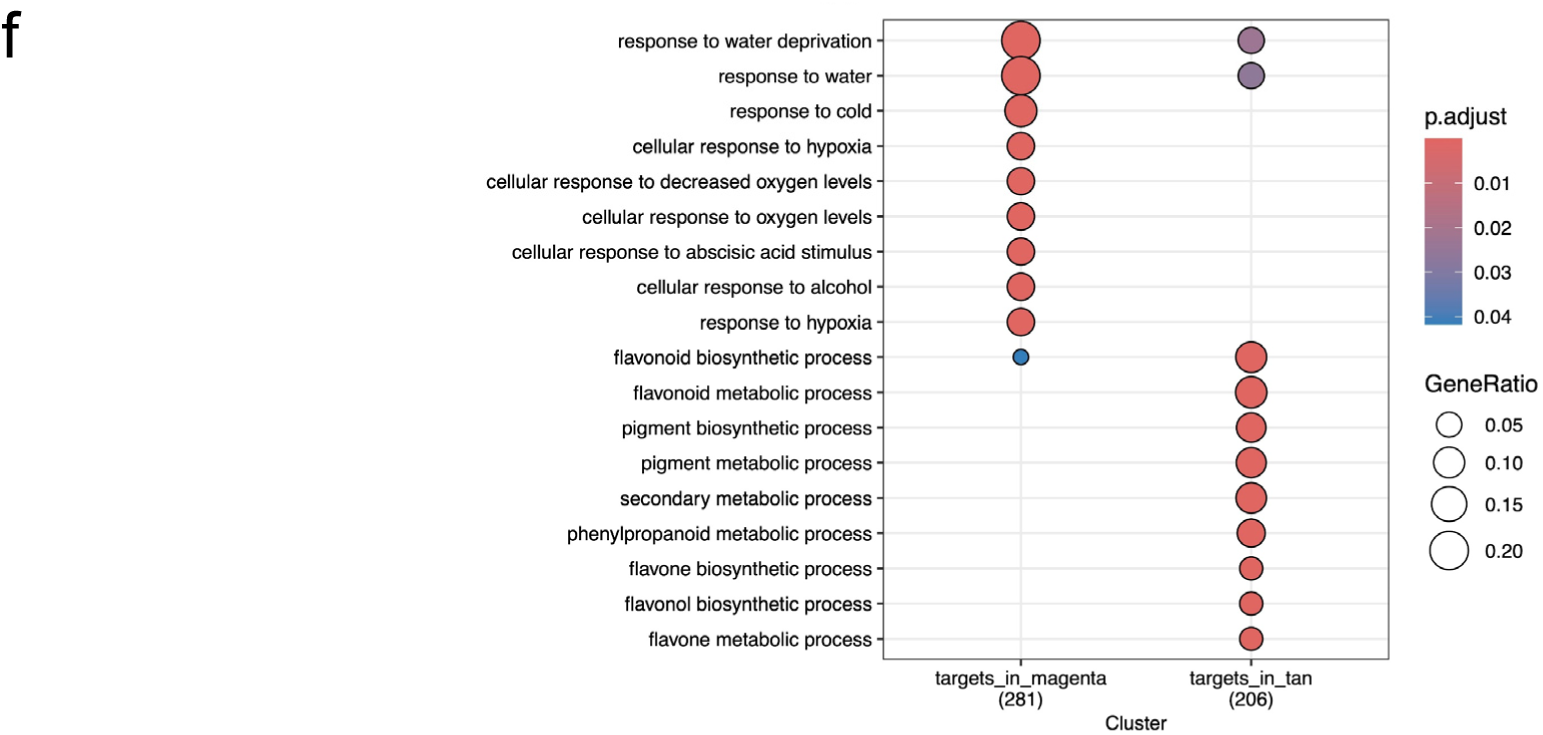
The predicted targets of the 40 potentially co-regulating TFs are associated with flavonoid processes (tan module) and with water, oxygen, and abscisic acid (ABA) processes (magenta module). The target genes in the tan and magenta co-expression modules displayed distinct GO terms, indicating that module membership was associated with differing biological processes. This confirms that our WGCNA network successfully grouped genes that are not only co-expressed, but also biologically correlated/coherent within each module. Indeed, the target genes in the tan module were significantly enriched for terms related to flavonoid and other metabolic processes, while those in the magenta module were significantly enriched for terms related to stress and environmental responses (water, cold, oxygen/hypoxia, and abscisic acid stimulus). This suggests that bHLH35 could be co-regulating genes associated with these two different biological signals. The four TFs with motifs significantly enriched in the blue module, and co-expressed with bHLH35, seem to target genes associated with heat and oxygen responses. In support of Fig. 4d.

**Fig. S8g.**
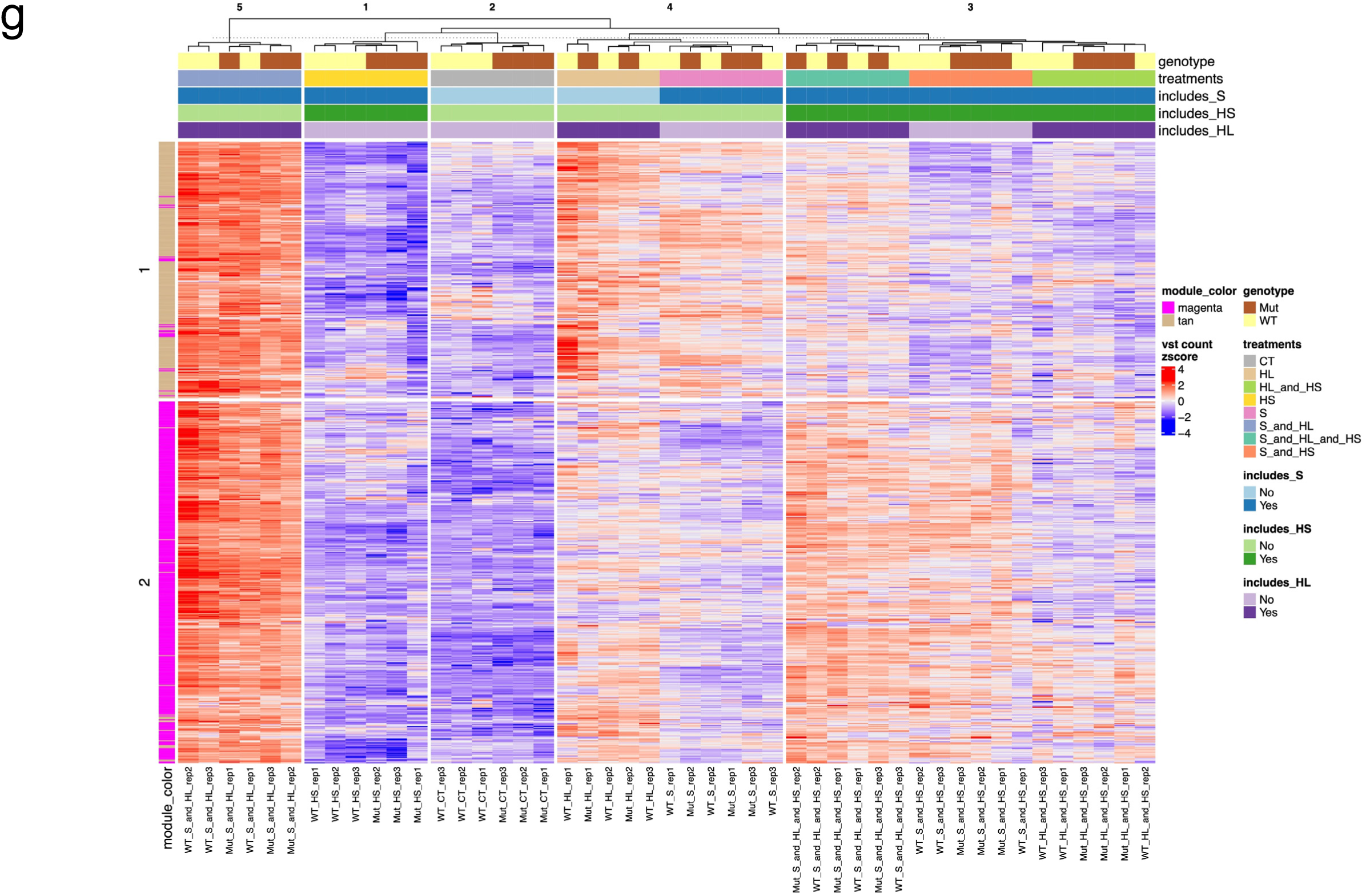
Expression patterns of the predicted target genes of the 40 TFs from the co-regulation model. The target genes in the tan and magenta modules follow a similar pattern where they are strongly upregulated (positive z-scores) in the ‘S and HL/EL’ treatment and strongly downregulated (negative z-scores) in the ‘HS’ and ‘CT’ treatments, regardless of genotype. The other combinations of treatments had weaker expression changes. Of note, the k-means clustering of the genes (rows in the heatmap) led to a grouping that are highly consistent with module membership, reaffirming the robustness of our WGCNA network. In support of Fig. 4d. Acronyms: HS = heat shock; HL = high light; S = salt.

**Fig. S9.**
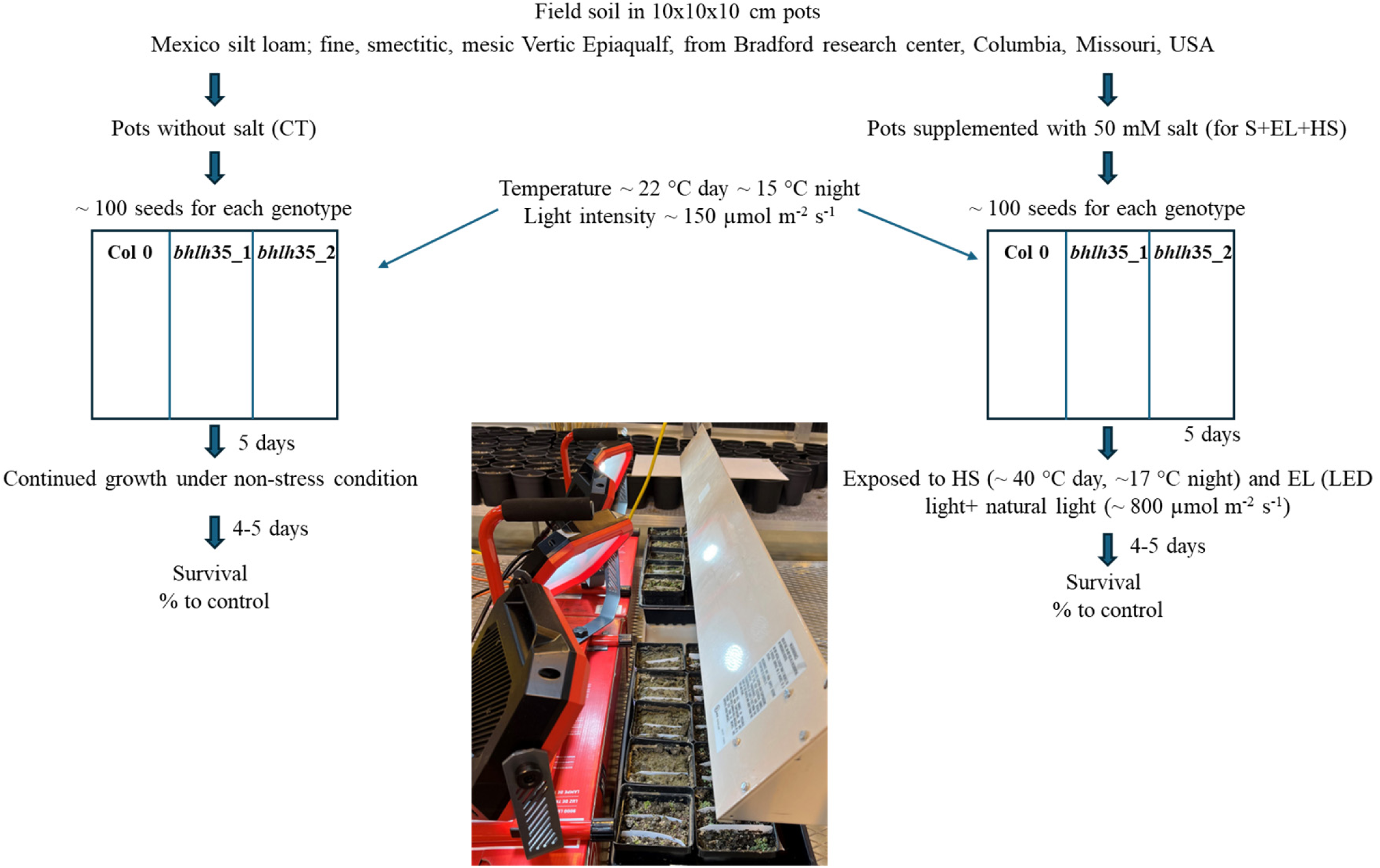
Experimental setup for simulated field conditions. Seeds of WT and the two *bhlh35* mutant lines were germinated side-by-side in plastic pots (10×10×10 cm) filled with field soil (Mexico silt loam; fine, smectitic, mesic Vertic Epiaqualf) obtained from the Bradford Research Center (Missouri Agricultural Experiment Station, Columbia, Missouri, USA, 38°53′N, 92°12′W). Soil for CT pots was watered with plain water, while soil for S+EL+HS was supplemented with 50 mM NaCl once at the beginning of the experiment. Seedlings were grown in a greenhouse (https://plantgrowthfacilities.missouri.edu/eastcampus.htm) under natural light shaded to 150 µmol m^−2^ s^−1^ at a temperature of ∼22 °C, day, ∼15 °C night. Five days post germination, CT non salt treated plants were kept under the conditions described above (CT conditions), while salt treated pots were subjected to EL and HS (for a combination of S+EL+HS), by removing them from shading, placing them under a heater (Kalglo®-Infrared Heater HS-2420 I 65” I 240V I 2000W, Fogelsville, PA, USA), and supplementing the natural daylight (500-600 µmol m^−2^ s^−1^) with LED lights (CRAFTSMAN 9000-Lumen LED twin light, model-CMXELAYMPL1029, CRAFTSMAN Mississauga, ON, CA; an additional 150-200 µmol µmol m^−2^ s^−1^ above natural conditions), for a total of 650-800 µmol m^−2^ s^−1^. Four to five days post stress application, the survival of CT and S+EL+HS treated plants was determined. In support of Fig. 1e.

### List of Supplementary Tables

**Table S1.** *Arabidopsis thaliana* loss- and gain-of-function mutants used in the study.

**Table S2.** Individual and combined stress treatment conditions used in the study.

**Table S3.** Transcripts significantly altered in their expression in wild type *Arabidopsis thaliana* plants subjected to S, EL, and/or HS stresses (in all possible combinations compared to control conditions).

**Table S4.** Transcripts significantly altered in their expression in *bhlh35 Arabidopsis thaliana* mutants subjected to S, EL, and/or HS stresses (in all possible combinations compared to control conditions).

**Table S5.** Transcripts with overlapping or unique expression included in Fig. 2a Venn diagrams.

**Table S6:** Transcripts with overlapping or unique expression included in Fig. 2b Upset plot.

**Table S7.** Class 1 transcripts (Figs. 2c, S4a).

**Table S8.** Class 2 transcripts (Figs. 2c, S4b).

**Table S9.** Description of genes in each node of the gene regulatory network analysis (Fig. S7).

**Table S10.** List of primers, probes, and promoter sequences used in the study.

**Table S11.** Description of genes in each node of the gene regulatory network analysis (Fig. S8).

